# Disruption of Prostaglandin F_2α_ Receptor Signaling Attenuates Fibrotic Remodeling and Alters Fibroblast Population Dynamics in A Preclinical Murine Model of Idiopathic Pulmonary Fibrosis

**DOI:** 10.1101/2023.06.07.543956

**Authors:** Luis R. Rodriguez, Soon Yew Tang, Willy Roque Barboza, Aditi Murthy, Yaniv Tomer, Tian-Quan Cai, Swati Iyer, Katrina Chavez, Ujjalkumar Subhash Das, Soumita Ghosh, Thalia Dimopoulos, Apoorva Babu, Caitlin Connelly, Garret A. FitzGerald, Michael F. Beers

**Affiliations:** Pulmonary, Allergy, and Critical Care Division Department of Medicine; Perelman School of Medicine at the University of Pennsylvania; Philadelphia, PA 19104; PENN-CHOP Lung Biology Institute; Perelman School of Medicine at the University of Pennsylvania; Philadelphia, PA 19104; Institute for Translational Medicine and Therapeutics; Department of Systems Pharmacology and Translational Therapeutics; Perelman School of Medicine at the University of Pennsylvania; Philadelphia, PA 19104; Calico Life Sciences LLC, South San Francisco, CA 94080

**Author notes:** These authors contributed equally to this work (Luis R. Rodriguez, Soon Yew Tang, Willy Roque Barboza).

**Keywords:** Idiopathic Pulmonary Fibrosis, Prostaglandin Signalling, Surfactant Biology, Adventitial Fibroblasts

## Abstract

Idiopathic Pulmonary Fibrosis (IPF) is a chronic parenchymal lung disease characterized by repetitive alveolar cell injury, myofibroblast proliferation, and excessive extracellular matrix deposition for which unmet need persists for effective therapeutics. The bioactive eicosanoid, prostaglandin F2*α*, and its cognate receptor FPr (*Ptfgr*) are implicated as a TGF*β*1 independent signaling hub for IPF. To assess this, we leveraged our published murine PF model (I^*ER*^ *−Sftpc*^*I*73*T*^) expressing a disease-associated missense mutation in the surfactant protein C (*Sftpc*) gene. Tamoxifen treated I^*ER*^ -*Sftpc* ^*I*73*T*^ mice develop an early multiphasic alveolitis and transition to spontaneous fibrotic remodeling by 28 days. I^*ER*^ -*Sftpc* ^*I*73*T*^ mice crossed to a Ptgfr null (FPr^*−*/*−*^) line showed attenuated weight loss and gene dosage dependent rescue of mortality compared to FPr^+/+^ cohorts. I^*ER*^ -*Sftpc* ^*I*73*T*^ /FPr^*−*/*−*^ mice also showed reductions in multiple fibrotic endpoints for which administration of nintedanib was not additive. Single cell RNA sequencing, pseudotime analysis, and in vitro assays demonstrated *Ptgfr* expression predominantly within adventitial fibroblasts which were reprogrammed to an “inflammatory/transitional” cell state in a PGF2*α*/ FPr dependent manner. Collectively, the findings provide evidence for a role for PGF2*α* signaling in IPF, mechanistically identify a susceptible fibroblast subpopulation, and establish a benchmark effect size for disruption of this pathway in mitigating fibrotic lung remodeling.

## Introduction

Idiopathic Pulmonary Fibrosis (IPF) is the most common subtype within a larger family of fibrosing parenchymal lung diseases in older adults (1, 2). The *sine qua non* IPF histology is the Usual Interstitial Pneumonitis (UIP) pattern composed of temporally and spatially heterogeneous areas of fibroblast/myofibroblast accumulation coupled with extracellular matrix deposition, disruption of alveolar architecture, and subpleural honeycombing (3-5). The typical clinical experience resulting from unchecked fibroproliferation of IPF patients is marked by progression of symptoms from cough and dyspnea to end-stage respiratory insufficiency resulting in lung transplantation or death within 3-5 years of diagnosis (6, 7). While two FDA approved anti-fibrotic agents (Pirfenidone; Nintedanib) emerged in 2014 (8, 9), the collective experience in IPF drug discovery and development over the past 30 years has been an overwhelming series of late failures in clinical trials highlighting the continued unmet need as well as an incomplete understanding of its complex underlying pathobiology (10, 11).

Although knowledge gaps in IPF pathogenesis persist, the past two decades have seen progress with a paradigm shift which offers hope for new IPF discovery. Consensus has pushed the molecular origins of IPF towards a pivotal role for alveolar epithelial cells as a primary upstream driver of aberrant injury and repair (12, 13). Specifically, the postulated contributions of alveolar type 2 (AT2) cell dysfunction along with disrupted alveolar niche cellular crosstalk to the development of a fibrotic lung phenotype has gained momentum wherein microfoci of repeated cycles of AT2 injury produce a dysfunctional repair process reminiscent of other fibrotic diseases observed in skin, kidney, and liver (14, 15). Throughout this evolution, there has been no doubt about the role of fibroblasts as the pathological extracellular matrix producing population(s). With the emergence of single cell RNA sequencing (scRNAseq), a complex heterogeneity in the fibroblast subpopulations that make up the IPF lung has been uncovered providing identification and classification of multiple *Col1a1*^+^ fibroblast subsets capable of participating in fibrogenic remodeling (16-23). Moreover, beyond the diseased environment, lung fibroblast heterogeneity is highly pertinent to homeostatic and developmental lung biology as the role for fibroblast subpopulations in direct support of the epithelium is clear (24-27). Thus, critical questions for IPF to be addressed include how a supportive mesenchyme enters a pathological state and what signaling pathways may promote this disruption.

Among the postulated mediators of crosstalk between these cellular components in the fibrotic niche, TGF*β* along with PDGF, CTGF, and FGFs have received major attention (11). While antagonizing TGF*β* has shown some therapeutic promise, off-target effects associated with blockade of its activation or signal transduction remain barriers to translation to the clinic (28). Similarly, in both preclinical models and clinical practice, either Pirfenidone or Nintedanib do not completely attenuate either fibrotic endpoints or loss of lung function suggesting additional pathways to fibrogenesis are contributors (8, 9, 29, 30).

Prostaglandins (PGs), secretory lipid mediators generated from arachidonic acid (AA), are multifaceted molecules that play critical roles in physiological balance, inflammation, and fibrosis (31). One prostaglandin in particular, Prostaglandin F2*α* (PGF2*α*), signaling through its cognate F-prostanoid receptor (FP receptor = FPr), has emerged as a potential facilitator of lung fibrogenesis. In a preclinical genetic model, the fibrotic response to exogenous bleomycin-induced injury was attenuated in Ptgfr (FPr) null mice (32). Furthermore, PGF2*α* metabolites are elevated in plasma of subjects with IPF (32, 33). While PGF2*α* has also been shown *in vitro* to prompt fibroblast proliferation and collagen production in a TGF*β*1 independent fashion, neither the effect size, specific PGF2*α* dependent fibroblast subpopulations, nor the integration of this signaling axis into other preclinical PF models or human IPF have been defined. Given the realities above, the complexities of IPF as a polycellular disease, and the many issues imparted by the use of exogenous injury models, we set out to define further the role of the PGF2*α* axis in fibrotic lung remodeling by employing our murine model of spontaneous epithelial-driven lung fibrosis (I^*ER*^-*Sftpc*^*I*73*T*^) shown both to recapitulate pathological and clinical features of the human IPF / UIP (34). Using both a genetic and pharmacologic approach, we defined a role for PGF2*α*/FPR signaling in lung fibrosis, demonstrating an effect size for FP antagonism that is non-inferior to that observed with nintedanib. Furthermore, scRNAseq analysis of the I^*ER*^-*Sftpc*^*I*73*T*^ model detected the emergence of a *Col1a1* fibroblast subpopulation which is dependent upon PGF2*α* / FPr signaling. Taken together, the results presented here implicate a role for PGF2*α* as a mediator of downstream events in the pathogenesis of IPF through modulation of fibrogenic programs in mesenchymal subpopulations.

## METHODS

### In vivo Mouse Models

*Mouse Model of Tamoxifen Induced Sftpc*^*I*73*T*^ *Expression* : Tamoxifen inducible *Sftpc*^*I*73*T*/*I*73*T*^ Rosa26ERT2FlpO^+/+^ (= I^*ER*^-*Sftpc*^*I*73*T*^) mice expressing an NH2-terminal HA-tagged murine *Sftpc*^*I*73*T*^ mutant allele into the endogenous mouse *Sftpc* locus were previously generated as reported (34) and are summarily detailed in Supplemental Methods. Tamoxifen treatment of adult I^*ER*^-*Sftpc*^*I*73*T*^ mice was initiated at 12-14 weeks of age by either intraperitoneal (ip) or oral gavage (OG) as indicated. Both male and female animals were used for the studies.

### *Generation of F Prostanoid receptor (FPr) deficient I*^*ER*^*-Sftpc*^*I*73*T*^ *mouse model*

Homozygous FPr knockout mice have been previously described (32) and were kindly provided by Shuh Narumiya (Kyoto University Faculty of Medicine, Japan). The breeding scheme to generate triple homozygous is detailed in *Supplemental Methods*. All mouse strains and genotypes generated for these studies were congenic with C57BL/6. Both male and female animals (aged 8-14 weeks) were utilized in tamoxifen induction protocols. All mice were housed under pathogen free conditions in an AALAC approved barrier facility at the Perelman School of Medicine, University of Pennsylvania. All experiments were approved by the Institutional Animal Care and Use Committee (IACUC) at the University of Pennsylvania.

### Reagents and Materials

Cytological stains used were Diff-Quik (Thermo Fisher Scientific, Inc., Pittsburgh, PA) and Giemsa (GS500; MilliporeSigma (St. Louis, MO)). Tamoxifen (non-pharmaceutical grade) was purchased from MilliporeSigma. Nintedanib was purchased from Cayman Chemical. OBE022 (35, 36) and BAY6872 (37) were manufactured for Calico LLC by Abbvie, Inc. Except where noted, all other reagents were electrophoretic or immunological grade and purchased from either MilliporeSigma or ThermoFisher. Antibodies used for flow-cytometry and FACS were obtained from commercial sources (**Supplemental Table 1**).

### Lung Histology

Whole lungs were fixed by tracheal instillation of 10% neutral buffered formalin (MilliporeSigma) at a constant pressure of 25 cm H_2_O. 6 *μ*M sections were stained with Hematoxylin&Eosin (H&E) or Masson’s Trichrome stains by the Pathology Core Laboratory of Children’s Hospital of Philadelphia. Slides were scanned using an Aperio ScanScope Model: CS2 (Leica) at 40X magnification and representative areas captured and exported as TIF files and processed in Adobe Illustrator

### Picrosirius Red (PSR) Staining

Staining of lung sections for fibrillar collagen was performed using the Picrosirius Red Stain Kit following the manufacturer’s instructions (Polysciences, Inc., Warrington PA). Digital morphometric measurements were performed on multiple lobes and multiple levels with ten random peripheral lung images devoid of large airways per slide analyzed at a final magnification of 100× using Image J as published (34, 38, 39). The mean area of each lung field in each section staining for PSR was calculated and expressed as a percentage of total section area as adapted from Henderson, *et al* (40).

### Bronchoalveolar Lavage Fluid (BALF) collection and processing

BALF collected from mice using sequential lavages of lungs with five X 1 ml aliquots of sterile saline was processed for analysis as described (41, 42). Cell pellets obtained by centrifuging BALF samples at 400 × g for 6 minutes were re-suspended in 1 ml of PBS, and total cell counts determined using a NucleoCounter (New Brunswick Scientific, Edison, NJ). Differential cell counts were determined manually from BALF cytospins stained with modified Giemsa (Sigma Aldrich, GS500) to identify macrophages, lymphocytes, eosinophils and neutrophils. Total protein content of cell free BALF was determined using the DC Protein Assay Kit (Cat 5000111; BioRAD, Inc, Hercules 3 Prostaglandin F2*α* Signaling in Lung Fibrosis CA) with bovine serum albumin as a standard according to the manufacturer’s instructions.

### Determination of BALF Soluble Collagen Content and TGFβ1 Concentration

Total acid soluble collagen content in cell free BALF was determined using the Sircol assay kit (Biocolor, Ltd; Carrickfergus, UK) according to the manufacturer’s instructions. TGF*β* concentration in the cell free BALF was calculated using Mouse TGF*β* 1 DuoSet Elisa (R&D Systems Cat DY1679-05) according to the manufacturer’s instructions.

### RNA Isolation and Quantitative Real Time Polymerase Chain Reaction

RNA was extracted from homogenized lung or isolated fibroblasts using RNeasy Mini Kit (Qiagen, Valencia, CA) following the manufacturer’s protocol. The concentration and quality of extracted RNA from the lung tissues were measured using NanoDrop One (Thermo Scientific, Wilmington, DE) and reverse-transcribed into cDNA using either Taqman Reverse Transcription Reagents (Applied Biosystems, Foster City, CA) or Verso cDNA Synthesis Kit (ThermoFisher). Quantitative real time PCR (qRT-PCR) for whole lung *Col1a1, Col1a2*, and *Ptgfr* was performed using Taqman Gene Expression Assays in an Applied Biosystems ViiA 7 real-time PCR system with a 384 well plate. Results were normalized to *Hprt*. Whole lung *Sftpc* as well as fibroblast lysate *Ces1d, Col13a1, Col14a1, Slc7a10, Pi16, Ebf1, Col14a1, Sfrp1, Hp, Tgfb1*, and *Cthrc1* were measured by qRT-PCR on a QuantStudio 7 Flex Real-Time PCR System with results normalized to *18S* and *Actb* RNA. Primer sequences for all mouse genes are listed in **Supplemental Table 1**.

### Single cell RNA sequencing and Analysis of Lung Cell Populations

To capture representative proportions of all major parenchymal and immune cell populations, we profiled 94,258 cells from the model, single cell suspension prepared by physical and enzymatic dissociation followed by MACS sorting by LS columns (Miltenyi Biotec 130-042-401) with CD45+ cell removal using CD45 micobeads (Miltenyi Biotec 130-052-301) followed by followed by “spike-back” of immune cells (20% of final suspensions) with biological replicates for each timepoint loaded onto individual GemCode instrument (10X Genomics), two for each of the timepoints. Single-cell barcoded droplets were produced using 10X Single Cell 3’ v3 chemistry. Libraries generated were sequenced using the HiSeq Rapid SBS kit, and the resulting libraries were sequenced across the two lanes of an Illumina HiSeq2500 instrument in a High Output mode. Single cell RNA-Seq reads were aligned to mouse genome (mm10/GRCm38) using STARSolo (version 2.7.5b). After initial quality control and processing, we analyzed the scRNA-seq data using the Scanpy pipeline (43). Genes expressed in fewer than 3 cells were removed, and cells with fewer than 200 genes and a mitochondrial fraction of less than 20% were excluded. Counts were log-normalized using `scanpy.pp.normalize_per_cell(counts_per_cell`=1*x*10^4^), followed by scanpy.pp.log1p.To integrate data from multiple samples, we used Scvi-tools (44). We applied `scvi.model.SCVI.setup_anndata` to establish the model parameters for integration, including: layer, `categorical_covariate_keys`, and `continuous_covariate_keys`. We then performed a principal component analysis (PCA) and generated a K-nearest neighbor (KNN) graph using scanpy.pp.neighbors with `n_neighbors=15`. The resulting KNN graph was used to perform Uniform Manifold Approximation and Projection (UMAP) dimension reduction to visualize the cells in two dimensions using scanpy.tl.umap(). Clustering was performed using the Leiden algorithm with scanpy.tl.leiden (45). We identified cell populations using known canonical marker genes or by assessing cluster-defining genes based on differential expressions. Additionally, we performed linear trajectory inference on the UMAP reduction using scFates (46) with the adventitial fibroblast cluster and the alveolar fibroblast cluster as the starting point and without assigned endpoints. Finally, we performed gene ontology analysis for enriched biological processes using GSEApy (47) based on differentially enriched genes between the groups.

### Multichannel Flow Cytometry For Identification of Lung Cell Populations

Flow cytometry was performed as we described (34, 39, 42, 48). Blood free perfused lungs were digested in Phosphate Buffered Saline (Mg and Ca free) with 2 mg/ml Collagenase Type I (Gibco Cat 17100017) and 50 units of DNase (Millipore Sigma Cat D5025), passed through 70/*mu*m nylon mesh to obtain single-cell suspensions, and then processed with ACK Lysis Buffer (Thermo Fisher). Cell pellets collected by centrifugation were resuspended in PBS+0.1% sodium azide and aliquots removed for determination of cell number using a NucleoCounter (New Brunswick Scientific, Edison, NJ). Cells were incubated with antibody mixtures (or isotype controls) and conjugated viability dye (see **Supplemental Table 1**). Stained cells were analyzed with an LSR Fortessa (BD Biociences). Cell populations were defined, gated, and analyzed with FlowJo software (FlowJo, LLC, Ashland, Oregon). Immune populations were identified by forward and side scatter followed by doublet discrimination of CD45+ viable cells and a sorting strategy modified from Misharin, *et al*. (49).

### Isolation, In vitro Culture, and Challenge of Mesenchymal Populations

Generation of single cell suspension and processing was performed as above. Mesenchymal populations were stained using a modified sorting strategy from Tsukui, *et al*. (20) with a BD FACSAria II (BD Biosciences) by the Flow Cytometry Core at the University of Pennsylvania. Collected cell populations were immediately seeded in tissue culture at a concentration of 2*x*10^5^ cells per 1.9 cm^2^ in DMEM containing 5% FBS and 10 ng/mL TGF*β* (Biolegend Cat 763102) or 500 nM Prostaglandin F (Cayman Chemical Cat 16010).

### Measurement of Urinary Prostanoids

Urinary prostanoid metabolites were measured by liquid chromatography/mass spectrometry as described (50). Such measurements provide a noninvasive, time integrated measurement of systemic prostanoid biosynthesis (51). Briefly, mouse urine samples were collected using metabolic cages over an eight hour period (9am to 5pm). Systemic production of PGI_2_, PGE_2_, PGD_2_, and TxA_2_ was determined by quantifying their major urinary metabolites-2, 3-dinor 6-keto PGF1*α* (PGIM), 7-hydroxy-5, 11-diketotetranorprostane-1, 16-dioic acid (PGEM), 11, 15-dioxo-9*α*-hydroxy-2, 3, 4, 5-tetranorprostan-1, 20-dioic acid (tetranor 4 Prostaglandin F2*α* Signaling in Lung Fibrosis PGDM) and 2, 3-dinor TxB_2_ (TxM), respectively. Results were normalized with urinary creatinine.

### Statistics

All data are presented with dot-plots and group mean ± SEM unless otherwise indicated. Statistical analyses were performed with GraphPad Prism (San Diego, CA). Student’s t-test (1 or 2 tailed as appropriate) were used for 2 groups; Multiple comparisons were done by analysis of variance (ANOVA) was performed with post hoc testing as indicated; survival analyses was performed using Kaplan Meier with Mantel Cox correction. In all cases statistical significance was considered at p values < 0.05.

## Data and code availability

The sequencing data generated in this study are deposited in Gene Expression Omnibus (GEO) (accession number will be released upon acceptance).

### Study Approval

Mice housed in pathogen free facilities were subjected to experimental protocols approved by the IACUC of the Perelman School of Medicine at the University of Pennsylvania.

## RESULTS

### Disruption of Prostaglandin F2α Signaling Reduces Morbidity, Mortality, and Fibrotic Endpoints in Sftpc^I73T^ Mice

As reported, the I^*ER*^-*Sftpc*^*I*73*T*^ knock-in mouse is a model of spontaneous lung fibrosis generated by “inducible” expression of the disease associated mutant SP-C^*I*73*T*^ protein from a single dose of intraperitoneally administered tamoxifen (TAM) (34). To assess the role of PGF2*α* signaling in pulmonary fibrosis, we first employed a genetic approach by crossing SP-C^*I*73*T*^ mice to a *Ptgfr* deficient line (**Figure S1**) with resultant lung *Ptgfr* mRNA levels reflecting the loss of one or two alleles (**Figure 1A**) and without changes in overall urinary prostanoid levels (**Figure S1 B**). Challenge of the resultant genotypes by induction of *Sftpc*^*I*73*T*^ was performed using tamoxifen dissolved in corn oil delivered either via oral gavage (Days 0,4) or by intraperitoneal (ip) injection (Day 0,3) (**Figure 1B**). Regardless of route of administration, each induction modality resulted in similar degrees of weight loss at 14 days (**Figure S2 B**) as well as comparable levels of mutant *Sftpc*^*I*73*T*^ gene expression (**Figure S2 C**) and BALF cell counts (**Figure S2 D**) at 28 days that were equivalent to single ip dosing of I^*ER*^-*Sftpc*^*I*73*T*^ /*Ptgfr*^+/+^ controls. However, TAM induction of I^*ER*^-*Sftpc*^*I*73*T*^ /*Ptgfr*^*−*/*−*^ resulted in decreased weight loss compared to I^*ER*^-*Sftpc*^*I*73*T*^ /*Ptgfr*^+/+^ controls (**Figure 1C**) and Kaplan-Meier analysis encompassing multiple cohorts demonstrated a *Ptgfr* allelic gene dose-dependent effect on survival (**Figure 1D**).

**Figure 1.**
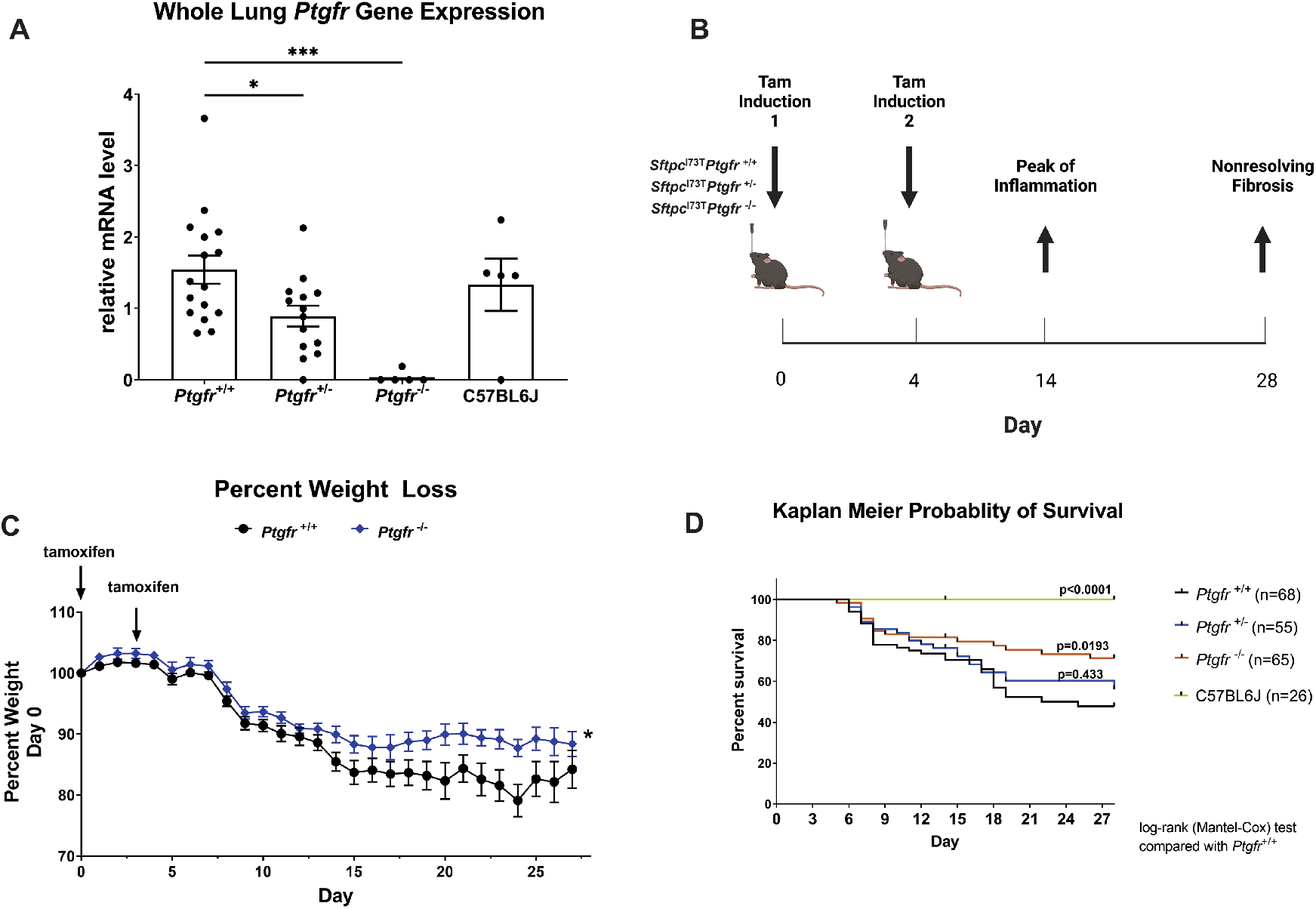
Deletion of Ptgfr reduces morbidity and mortality in I^ER^-Sftpc^I73T^ mice. (A) *Ptgfr* mRNA content of whole lung mRNA isolated from generated lines of I^*ER*^ -*Sftpc*^*I*73*T*^ /*Ptgfr* mice deficient in 0, 1, 2 *Ptgfr* alleles assayed by qRT-PCR; (B) Schematic of single, split ip or split OG dosing strategy employed for Tamoxifen induction of I^*ER*^ -*Sftpc*^*I*73*T*^ /*Ptgfr*^*−*/*−*^ and I^*ER*^ -*Sftpc*^*I*73*T*^ /*Ptgfr*^+/+^ cohorts; (C) Representative weight loss curve from a single cohort containing I^*ER*^ -*Sftpc*^*I*73*T*^ /*Ptgfr*^*−*/*−*^ (n=20) and I^*ER*^ -*Sftpc*^*I*73*T*^ /*Ptgfr*^+/+^(n=13) controls * P < 0.05 vs controls (D) Aggregate Kaplan Meier curve for I^*ER*^ -*Sftpc*^*I*73*T*^ /*Ptgfr*^*−*/*−*^ mice from 3 cohorts separately induced with either single ip or split ip doses of Tamoxifen in Corn Oil with total numbers of each Ptgfr genotype shown. Negative Controls consisted of *Sftpc*^*WT*^ C57BL/6 mice given Tamoxifen or uninduced I^*ER*^ -*Sftpc*^*I*73*T*^ /*Ptgfr*^*−*/*−*^ animals. p values versus I^*ER*^ -*Sftpc*^*I*73*T*^ /*Ptgfr*^*−*/*−*^ obtained by log-rank testing are shown.

I^*ER*^-*Sftpc*^*I*73*T*^ /*Ptgfr* null animals were significantly protected from *Sftpc*^*I*73*T*^ induced pulmonary fibrosis. Lung sections prepared from Day 28 animals showed marked changes in lung Trichrome collagen staining (**Figure 2A**) that were accompanied by quantitative reductions in collagen gene expression, BALF soluble collagen, and fibrillar collagen staining of lung sections. (**Figure 2B-D**).

**Figure 2.**
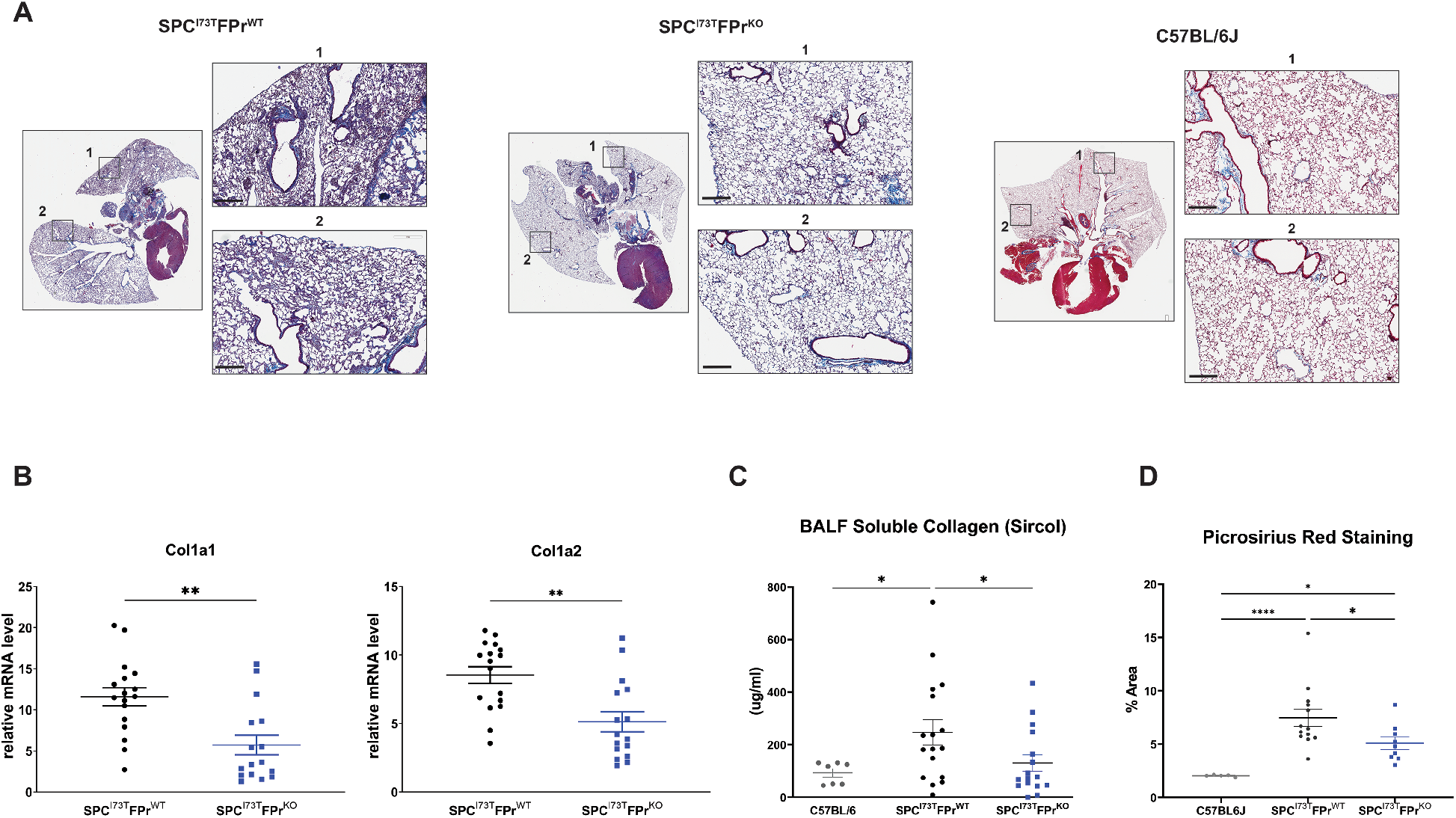
Ptgfr Deficiency Mitigates Collagen Expression and Deposition Post Induction of Sftpc^I73T^. (A)Representative histology from I^*ER*^ - *Sftpc*^*I*73*T*^ /*Ptgfr*^+/+^ and I^*ER*^ -*Sftpc*^*I*73*T*^ /*Ptgfr*^*−*/*−*^ mice 28 days after tamoxifen and development of fibrosis. Images are derived from Masson’s Trichrome stained sections, scale bars 300 *μ*M; (B) Relative fold mRNA levels between I^*ER*^ -*Sftpc*^*I*73*T*^ /*Ptgfr*^+/+^ and I^*ER*^ -*Sftpc*^*I*73*T*^ /*Ptgfr*^*−*/*−*^ measured via qPCR demonstrates decreased *Col1a1* and *Col1a2* in I^*ER*^ -*Sftpc*^*I*73*T*^ /*Ptgfr*^*−*/*−*^ mice 28 days after tamoxifen induction; (C) Quantification of soluble collagen in BALF from mice during fibrotic remodeling reveals a lower concentration in I^*ER*^ -*Sftpc*^*I*73*T*^ /*Ptgfr*^*−*/*−*^ mice; (D) Picrosirius red staining for collagen fibrils indicates mitigation of collagen deposition in I^*ER*^ -*Sftpc*^*I*73*T*^ /*Ptgfr*^*−*/*−*^ mice. Quantification performed using ImageJ, data represents percentage of total section area. All quantified data in this figure is derived from I^*ER*^ -*Sftpc*^*I*73*T*^ /*Ptgfr*^+/+^ (n=17) and I^*ER*^ -*Sftpc*^*I*73*T*^ /*Ptgfr*^*−*/*−*^ (n=16) *p < 0.05 ** p<0.005

The effect size of these changes was similar to that observed using the FDA approved antifibrotic Nintedanib in the same *Sftpc*^*I*73*T*^ preclinical model (**Figure S3**). Following oral TAM induction, mice were randomized on Day 12 to receive either Nintedanib or vehicle in a “treatment intervention” protocol (**Figure S3 A**). By Day 28, Nintedanib treatment resulted in significant reductions in BALF total protein, BALF soluble collagen, and lung fibrillar collagen as assessed by picrosirius red staining (**Figure S3 D, E, F**) accompanied by improvement in lung histology (**Figure S3 G**). The magnitude of Nintedanib mediated changes in these endpoints was consistent with prior published data in Bleomycin injured mice (30) as were the observed minor improvements in weight loss, BALF cell counts, and restrictive lung physiology (**Figure S3 B, C, H**).

### Nintedanib is Not Additive to FPr Signaling Inhibition in Modulating Fibrotic Endpoints

We next tested the combined efficacy of Nintedanib and FPr signaling inhibition in our genetic model. Using a 3armed protocol, cohorts of I^*ER*^-*Sftpc*^*I*73*T*^ /*Ptgfr*^+/+^ and I^*ER*^- *Sftpc*^*I*73*T*^ /*Ptgfr*^*−*/*−*^ were first induced with oral TAM and on Day 12 the I^*ER*^-*Sftpc*^*I*73*T*^ /*Ptgfr*^*−*/*−*^ cohort was randomized to receive either Nintedanib or vehicle (**Figure 3A**). As expected I^*ER*^-*Sftpc*^*I*73*T*^ /*Ptgfr*^*−*/*−*^ null mice were protected from weight loss and late mortality (**Figure 3B-C**) compared with I^*ER*^-*Sftpc*^*I*73*T*^ /*Ptgfr*^+/+^ animals, however, addition of Nintedanib provided no added improvement in morbidity, mortality, or any fibrotic endpoint including BALF soluble collagen (**Figure 3D**), fibrillar collagen deposition (**Figure 3E**), or lung histology (**Figure 3F**).

**Figure 3.**
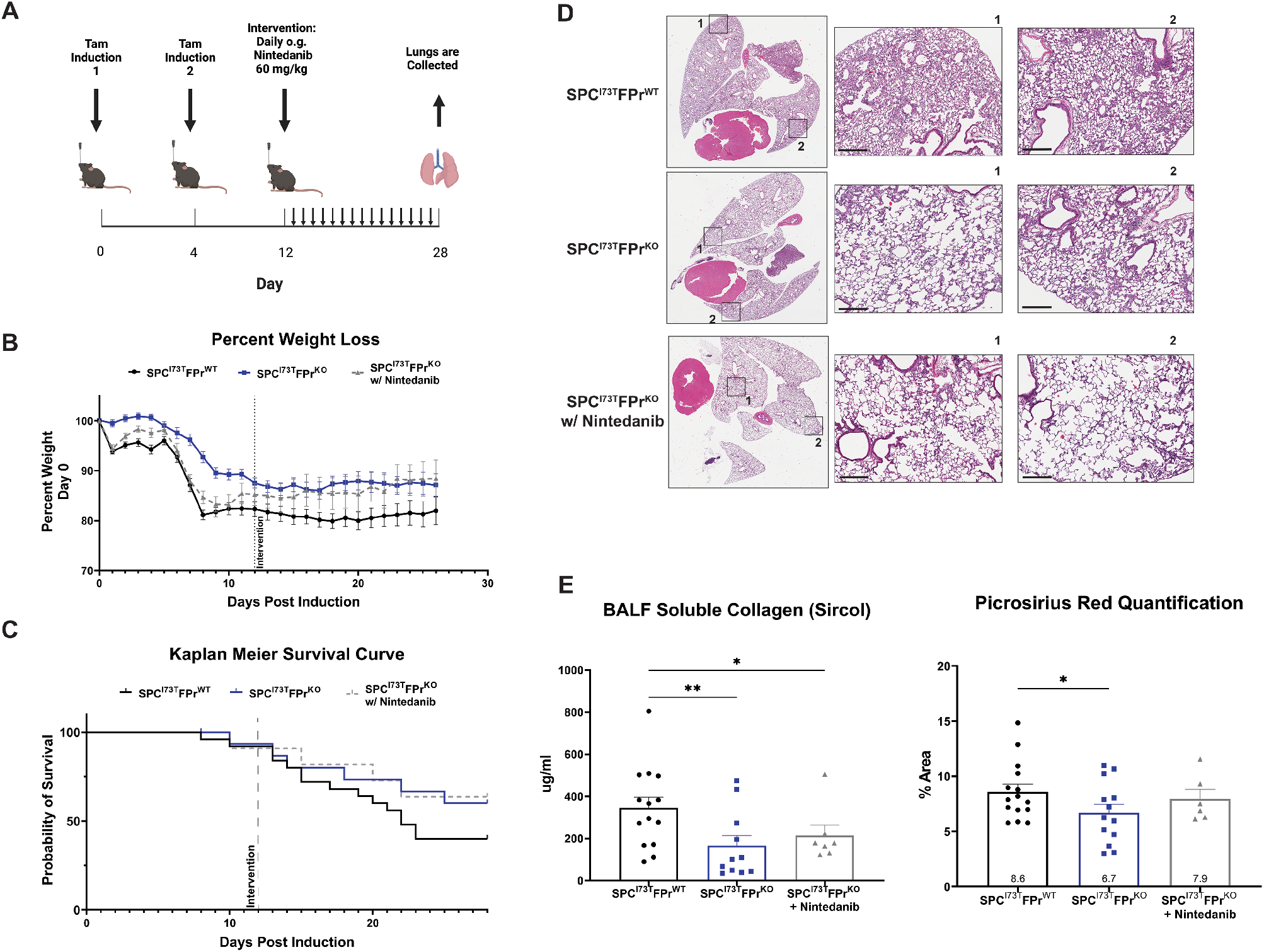
Nintedanib Intervention is not additive to Ptgfr Deficiency in I^ER^-Sftpc^I73T^ mice. (A) Daily nintedanib intervention (60mg/kg) was initiated at day 12 after tamoxifen induction. Following 16 days of intervention, surviving mice were euthanized and processed to evaluate fibrotic endpoints. (B) Weight loss as a percent of starting weight was tracked throughout the study, nintedanib intervention in I^*ER*^ -*Sftpc*^*I*73*T*^ /*Ptgfr*^*−*/*−*^ mice did not reduce mean weight loss (C) Kaplan Meier survival analysis demonstrates a non-significant improved probability of survival in I^*ER*^ - *Sftpc*^*I*73*T*^ /*Ptgfr*^*−*/*−*^, which was not improved through nintedanib intervention (D) Representative histology fromI^*ER*^ -*Sftpc*^*I*73*T*^ /*Ptgfr*^+/+^, I^*ER*^ - *Sftpc*^*I*73*T*^ /*Ptgfr*^*−*/*−*^, and nintedanib treated I^*ER*^ -*Sftpc*^*I*73*T*^ /*Ptgfr*^*−*/*−*^ mice 28 days after tamoxifen and development of fibrosis. Images are derived from H&E stained sections, scale bars 300 *μ*M; (E) Soluble collagen in BALF as measured by Sircol assay and fibrillar collagen in histological sections measured by PSR staining demonstrated a significant decrease in I^*ER*^ -*Sftpc*^*I*73*T*^ /*Ptgfr*^*−*/*−*^ mice, again nintedanib treatment did not improve these outcomes. Quantification of PSR was performed using ImageJ, data represents percentage of total section area. Survival and weight loss data is derived from I^*ER*^ -*Sftpc*^*I*73*T*^ /*Ptgfr*^+/+^ (n=26), I^*ER*^ -*Sftpc*^*I*73*T*^ /*Ptgfr*^*−*/*−*^ w/o nintedanib (n=12), and I^*ER*^ -*Sftpc*^*I*73*T*^ /*Ptgfr*^*−*/*−*^ w/ nintedanib (n=12). Soluble collagen and PSR analysis included I^*ER*^ -*Sftpc*^*I*73*T*^ /*Ptgfr*^+/+^ (n=14), I^*ER*^ -*Sftpc*^*I*73*T*^ /*Ptgfr*^*−*/*−*^ w/o nintedanib (n=11), and I^*ER*^ -*Sftpc*^*I*73*T*^ /*Ptgfr*^*−*/*−*^ w/ nintedanib (n=7) *p < 0.05 ** p<0.005

To corroborate these data sets, the protective effect of PGF2*α* signaling on lung fibrogenesis was also assessed pharmacologically using FPr antagonist tool compounds in a model of bleomycin-induced lung fibrosis (**Figure S4**). Both OBE022 and BAY6672 have been shown to exhibit potent and selective blockade of FPr receptor mediated signaling (35, 37). Using an intervention protocol, mice challenged with intratracheal bleomycin were randomized to receive drug or vehicle beginning on day six and animals taken down on Day 22 (**Figure S4 A**). In a study focused on histological endpoints, BAY6672 delivered twice daily (30 mpk; 100 mpk) produced significant reductions in collagen staining, Ashcroft scoring, and *α*-SMA staining at magnitudes like Nintedanib (**Figure S4 B**). In a separate experiment using OBE022, significant changes in body weight were not observed at each of two doses (100 and 300mpk). However, reductions in BALF soluble collagen and cell counts similar in magnitude to Nintedanib were observed while reductions in histological fibrosis scores were inferior to Nintedanib (**Figure S4 C**).

Taken together, these data are consistent with the hypothesis that FPr blockade produces an antifibrotic effect size similar to but not additive to Nintedanib.

### Ptgfr Deficiency Had no Effect on Early Lung Inflammation Induced by SP-C^I73T^

To exclude the possibility that the observed antifibrotic effect of FPr signaling blockade were related to upstream effects on lung inflammation/injury, we surveyed the I^*ER*^-*Sftpc*^*I*73*T*^ /*Ptgfr* model 14 days post-tamoxifen (the published peak of early inflammatory phase). As shown in **Figure 4**, we found that the absence of *Ptgfr* signaling produced no reduction in either BALF protein or total cell counts (**Figure 4A-B**). Differential counting of BALF cytospins revealed no alterations in the distribution of monocytes, eosinophils, neutrophils, or lymphocytes (**Figure 4C-D**). We also assessed immune populations in the lung parenchyma using flow cytometry analysis using a previously published protocol illustrated in **Figure S5A** and **S5C** ((34, 48, 52). When compared with *Ptgfr*^+/+^ controls, at Day 14, the induced I^*ER*^-*Sftpc*^*I*73*T*^ /*Ptgfr*^*−*/*−*^ cohort had similar levels of both myeloid and lymphocyte lineages (**Figure S5B**).

**Figure 4.**
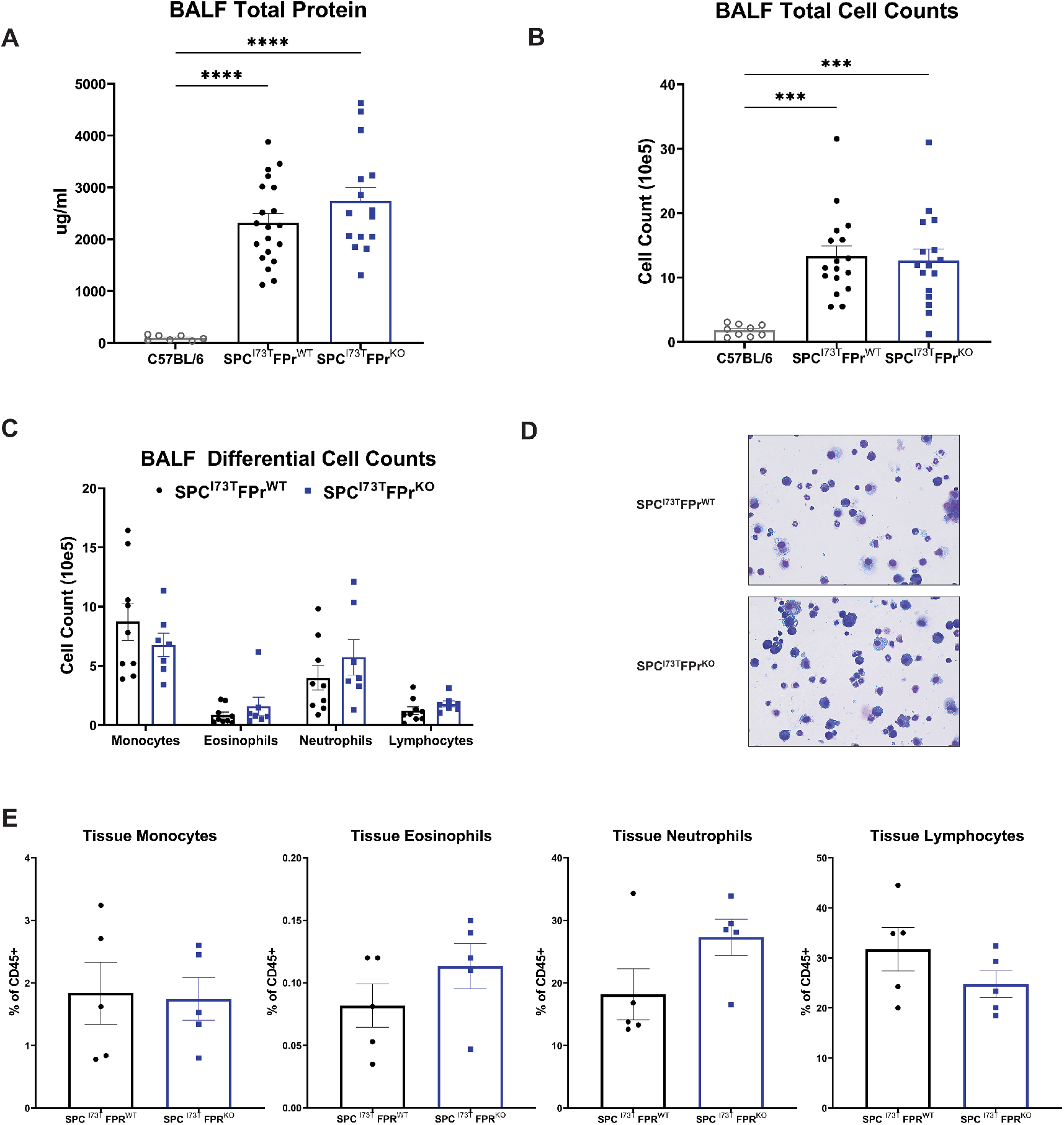
Ptgfr Deficiency Has No Effect on Early Lung Injury and Inflammation in I^ER^-Sftpc^I73T^. (A-B) Quantification of total protein and total cell counts in BALF did not result in a significant difference between I^*ER*^ -*Sftpc*^*I*73*T*^ /*Ptgfr*^+/+^ (n=17) and I^*ER*^ -*Sftpc*^*I*73*T*^ /*Ptgfr*^*−*/*−*^ (n=16) mice 14 days post tamoxifen induction. *** p < 0.005 **** p<0.0005 (C) BALF cell differential determined by quantification of modified Giemsa stained cytospins yielded no significant difference between I^*ER*^ -*Sftpc*^*I*73*T*^ /*Ptgfr*^+/+^ (n=9) and I^*ER*^ -*Sftpc*^*I*73*T*^ /*Ptgfr*^*−*/*−*^ (n=7) mice 14 days post tamoxifen induction. (D) Representative Giemsa stained images from I^*ER*^ -*Sftpc*^*I*73*T*^ /*Ptgfr*^+/+^ and I^*ER*^ -*Sftpc*^*I*73*T*^ /*Ptgfr*^*−*/*−*^ mice 14 days post tamoxifen induction. (E) Flow cytometry quantification of whole lung single cell suspensions confirms that there is no differential immune cell infiltration in between I^*ER*^ -*Sftpc*^*I*73*T*^ /*Ptgfr*^+/+^ (n=5) and I^*ER*^ -*Sftpc*^*I*73*T*^ /*Ptgfr*^*−*/*−*^ (n=5) mice 14 days post tamoxifen induction.

### Single Cell RNAseq Analysis Identifies Increased Ptgfr Expression in Adventitial Fibroblasts

A prior study has demonstrated that PGF2*α* treatment modifies fibroblast phenotypes *in vitro* (32). Given the immense heterogeneity within lung stromal cells at homeostasis and the dynamic changes in both surface markers and transcriptomic profiles that occur during fibrogenesis in other *in vivo* model systems (e.g. bleomycin) and human IPF (19, 21, 23, 24, 26, 27, 53, 54), we next performed single cell RNA sequencing (scRNAseq) on digested lung samples to assess transcriptional differences in specific mesenchymal subsets of the I^*ER*^-*Sftpc*^*I*73*T*^ model in the presence and absence of PGF2*α*-*Ptgfr* signaling (**Figure 5**). We induced cohorts of I^*ER*^-*Sftpc*^*I*73*T*^ /*Ptgfr*^+/+^ and I^*ER*^-*Sftpc*^*I*73*T*^ /*Ptgfr*^*−*/*−*^ animals with oral TAM and harvested the lungs at 14 and 28 days (n=2 per genotype per time point). Controls (n= 4) consisted of one mouse each of uninduced I^*ER*^-*Sftpc*^*I*73*T*^ /*Ptgfr*^+/+^, I^*ER*^-*Sftpc*^*I*73*T*^ /*Ptgfr*^*−*/*−*^, *Sftpc*^*WT*^ /*Ptgfr*^+/+^, and *Sftpc*^*WT*^ /*Ptgfr*^*−*/*−*^ genotypes. To balance stromal and effector cell numbers, resultant single cell suspensions were initially depleted using *α*-CD45 magnetic beads and captured CD45 cells then “spiked back” CD45 cells to achieve approximately 20% of the total cell numbers.

**Figure 5.**
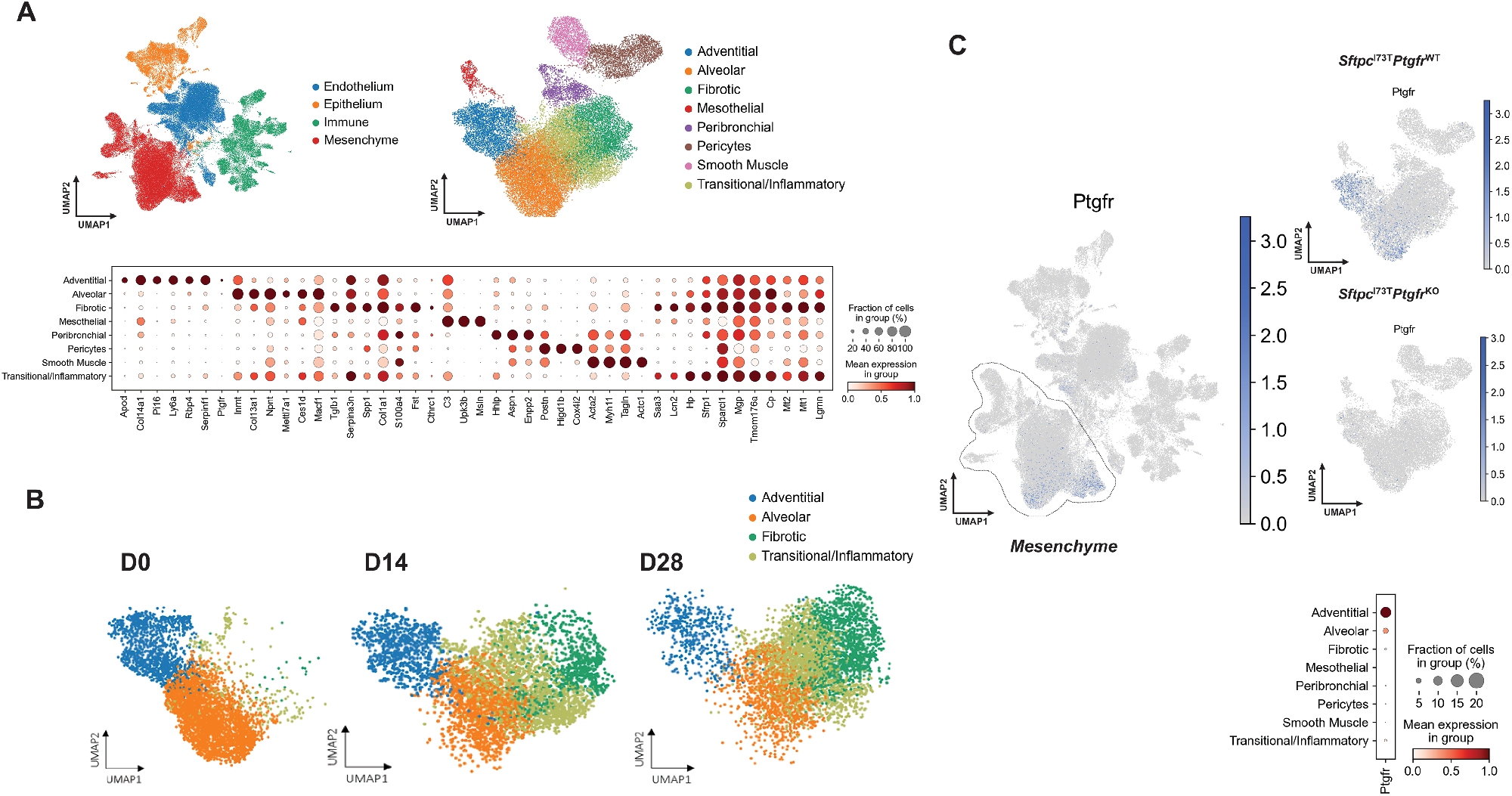
Ptgfr Expression is Limited to Adventitial and Alveolar Fibroblasts. (A) UMAP clustering 94258 cells identifies 4 primary cell compartments in I^*ER*^ -*Sftpc*^*I*73*T*^ /*Ptgfr*^+/+^, I^*ER*^ -*Sftpc*^*I*73*T*^ /*Ptgfr*^*−*/*−*^, and uninduced controls. Subclustering of the mesenchymal compartment identifies 8 mesenchymal clusters defined by marker genes depicted as a gradient dot plot. (B) UMAP projections of *Pdgfra*^+^ mesenchymal populations across time identifies two injury specific clusters (Fibrotic and Transitional/Inflammatory). (C) UMAP projection of all cells identifies the restriction of *Ptgfr* expression to the mesenchymal compartment and lack of Ptgfr expression in I^*ER*^ -*Sftpc*^*I*73*T*^ /*Ptgfr*^+/+^ mice. Gradient dot plot of *Ptgfr* expression within the mesenchyme demonstrates increased expression and increased percent expression of *Ptgfr* in adventitial fibroblasts as compared to alveolar fibroblasts

We profiled 35,002 cells from I^*ER*^-*Sftpc*^*I*73*T*^ /*Ptgfr*^+/+^, 32,594 cells from I^*ER*^-*Sftpc*^*I*73*T*^ /*Ptgfr*^*−*/*−*^, and 26,662 cells from pooled control animals and identified all major populations including endothelial, epithelial, mesenchymal, and effector (**Figure 5A and S6**). Removing non-mesenchymal cell (MC) populations defined using marker genes from multiple published studies and publicly available gene sets (**Figure S6**), we observed 8 well-segregated clusters of MC which each expressed a unique profile of marker genes (**Figure 5A**). These included alveolar fibroblasts or lipofibroblasts (*Npnt, Inmt, Ces1d*), adventitial fibroblast (*Col14a, Pi16, Apod*), fibrotic fibroblast (*Col1a1*^*High*^,*Tgfb1, Spp1, Cthrc1*), and a “transitional/inflammatory” population (*Sfrp1, Lcn2, Saa3, Hp*) recently defined by several groups (19-23). When stratified by time from induction of I^*ER*^-*Sftpc*^*I*73*T*^ model, most MC at Day 0 (uninduced or control) were in the alveolar and adventitial clusters. TAM induction results in time dependent increases in transitional/inflammatory and fibrotic populations commensurate with a loss of alveolar and adventitial cells (**Figure 5B**). Importantly, among these 8 MC populations *Ptgfr* expression predominantly localized in the adventitial fibroblast population with a lesser degree in alveolar cluster of *Ptgfr* genotypes (**Figure 5C**). We also failed to detect significant *Ptgfr* expression in endothelial, epithelial or immune cell clusters (**Figure S7**).

### Reprogramming of Adventitial Fibroblasts to the Transition/Inflammatory State is Ptgfr Dependent

The restriction of *Ptgfr* expression to the mesenchymal compartment, specifically adventitial and alveolar fibroblasts (**Figure 5C**), facilitated an assessment of the role of PGF2*α* signaling in lung fibroblast populations during the evolution of pulmonary fibrosis. We performed pseudotime analysis using scFates originating with either alveolar or adventitial populations and observed marked differences in *Ptgfr* dependence among these 2 populations to identify the potential progenitors of the fibrotic population (**Figure 6A-B**). Vectors originating from alveolar fibroblasts suggest entry into the transitional population prior to terminating in the fibrotic cluster and were unaffected by the loss of *Ptgfr*. (**Figure 6A**). In contrast, while I^*ER*^-*Sftpc*^*I*73*T*^ /*Ptgfr*^+/+^ adventitial fibroblasts were shown to be candidate progenitors for terminal fibrotic fibroblasts, the *Ptgfr*^*−*/*−*^adventitial population was markedly disrupted with the terminal node of this vector predicted within the transitional cluster suggesting a failure to progress (**Figure 6B**). Supporting the pseudotime analysis, gene expression profiles of the resulting fibrotic gene clusters from I^*ER*^-*Sftpc*^*I*73*T*^ /*Ptgfr*^+/+^, I^*ER*^-*Sftpc*^*I*73*T*^ /*Ptgfr*^*−*/*−*^, and *Sftpc*^*WT*^ /*Ptgfr*^+/+^ populations at Day 28 demonstrated that I^*ER*^-*Sftpc*^*I*73*T*^ /*Ptgfr*^*−*/*−*^ fibrotic clusters were relatively deficient in the fibrotic signature genes while retaining increased levels of transitional genes suggesting that a maximal fibrogenic response required intact PGF2*α* signaling (**Figure 6C**). Gene set enrichment analysis comparing the fibrotic clusters displayed a transcriptomic profile consistent with an attenuated fibrotic signature and decreased TGF*β* signaling in FPr KO mice (**Figure 6D**). Finally, analysis of TGF-*β*1 in BALF identifies a *Ptgfr* dependent decrease in TGF*β*1, confirming the transcriptomic analysis and suggesting that terminal fibrotic fibroblasts are a primary source of TGF*β*1 in the *Sftpc*^*I*73*T*^ model.

**Figure 6.**
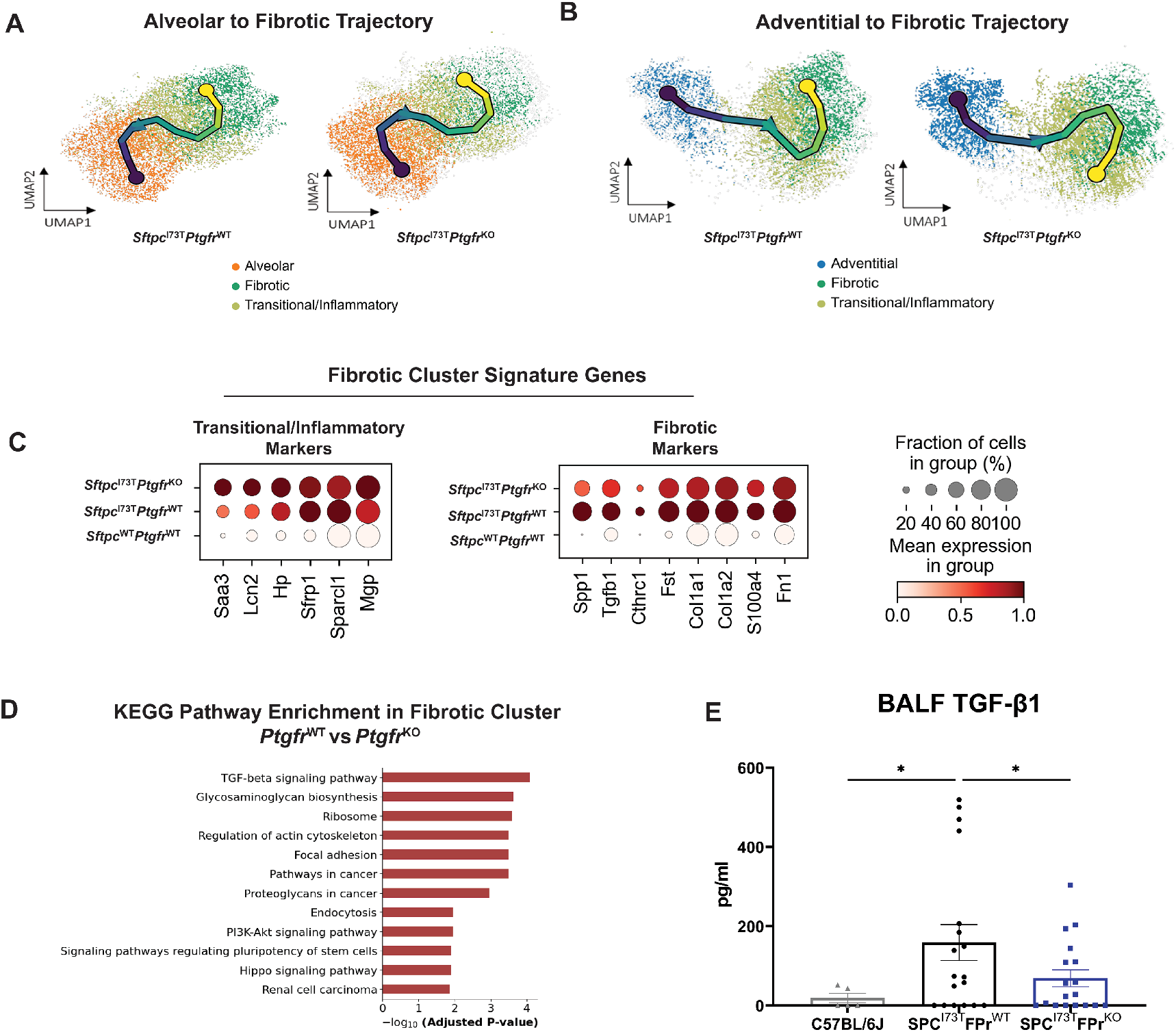
Ptgfr Deficiency Alters Fibroblast Lineage Trajectory Through Fibrotic Remodeling. A) UMAP analysis of reclustered alveolar, transitional/inflammatory, and fibrotic fibroblasts with superimposed vector from pseudotime trajectory analysis reveals no *Ptgfr* dependent effect on terminal node. (B) UMAP analysis of reclustered adventitial, transitional/inflammatory, and fibrotic fibroblasts with superimposed vector from pseudotime trajectory analysis demonstrates a *Ptgfr* dependent effect on terminal node. In I^*ER*^ -*Sftpc*^*I*73*T*^ /*Ptgfr*^*−*/*−*^ the terminal node is found in the transitional/inflammatory cluster while I^*ER*^ -*Sftpc*^*I*73*T*^ /*Ptgfr*^+/+^ samples have a vector terminating in the fibrotic cluster. (C) Comparative analysis of gene expression within the fibrotic cluster of I^*ER*^ -*Sftpc*^*I*73*T*^ /*Ptgfr*^+/+^ and I^*ER*^ -*Sftpc*^*I*73*T*^ /*Ptgfr*^*−*/*−*^ mice is presented by gradient gene expression dot plots. Marker genes associated with the transitional/inflammatory cluster are comparatively elevated in I^*ER*^ -*Sftpc*^*I*73*T*^ /*Ptgfr*^*−*/*−*^ mice while fibrotic marker genes are elevated in the I^*ER*^ -*Sftpc*^*I*73*T*^ /*Ptgfr*^+/+^ mice. (D) KEGG pathway enrichment analysis comparing the fibrotic clusters identifies multiple pathways associated with cytoskeletal rearrangement, mesenchymal activation, and TGF*β* signaling that are upregulated in I^*ER*^ -*Sftpc*^*I*73*T*^ /*Ptgfr*^+/+^ mice. (E) Measurement of BALF TGF*β*1 via elisa demonstrates a significant decrease in I^*ER*^ -*Sftpc*^*I*73*T*^ /*Ptgfr*^*−*/*−*^ mice (n=18) as compared to I^*ER*^ -*Sftpc*^*I*73*T*^ /*Ptgfr*^+/+^ mice (n=18). *p < 0.05

To validate the computational analysis, we next assessed the ability of PGF2*α* signaling to promote adventitial entry into the transitional state *in vitro* (**Figure 7**). Employing cell surface markers identified in prior published studies (19-21), we first isolated adventitial and alveolar populations from bulk mesenchyme prepared from *Sftpc*^*WT*^ /*Ptgfr*^+/+^ and *Sftpc*^*WT*^ /*Ptgfr*^*−*/*−*^ mice using FACS (**Figure 7A**). When analyzed by qPCR, the resultant mesenchymal populations (alveolar and adventitial) were each enriched in population-specific marker genes (**Figure 7B**). Consistent with the scRNAseq derived bioinformatic predictions, after culture and challenge with PGF2*α*, adventitial but not alveolar fibroblasts acquired markers of the transition state (*Sfrp1; Hp*) that was FPr dependent (**Figure 7C**). Notably, PGF2*α* treatment of either population failed to stimulate markers associated with the fibrotic myofibroblast cluster while both adventitial and alveolar fibroblast populations significantly increased pathologic fibrotic marker expression (*Cthrc1; Col1a1*) in response to TGF*β* in a manner independent of FPr status. Interestingly, commensurate with these changes, TGF*β* also downregulated markers associated with the transition/inflammatory state (**Figure 7D**). These data further support a role for PGF2*α*/ FPr mediated signaling in selectively modulating the transcriptomic trajectory of an important MC progenitor population capable of contributing to fibrogenesis.

**Figure 7.**
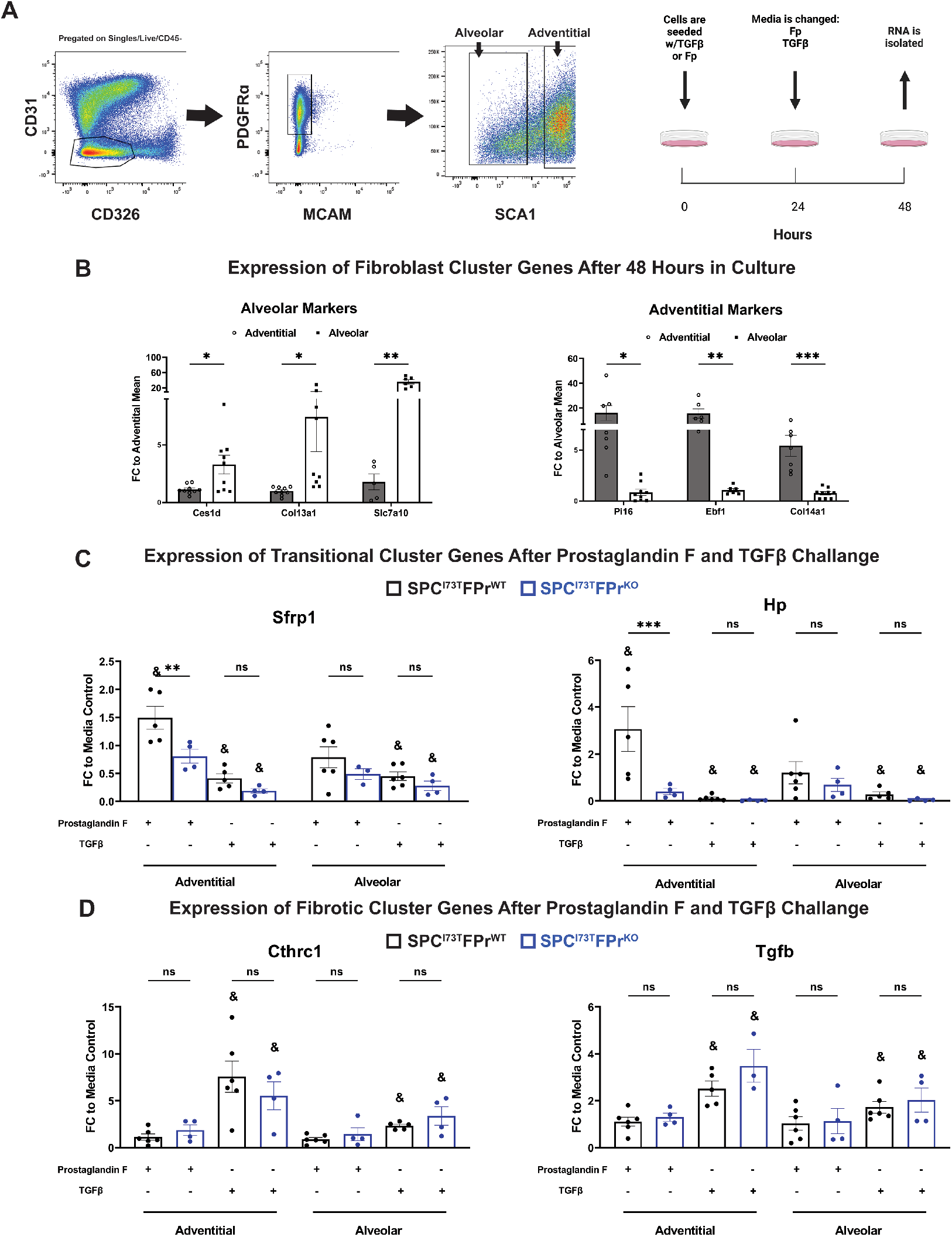
In vitro Prostaglandin F2α (Fp) Challenge Promotes Adventitial Fibroblast Entry Into the Transitional/Inflammatory State. (A) Sorting strategy for the isolation of adventitial and alveolar fibroblast used in I^*ER*^ -*Sftpc*^*I*73*T*^ /*Ptgfr*^+/+^ and I^*ER*^ -*Sftpc*^*I*73*T*^ /*Ptgfr*^*−*/*−*^ prior to induction by tamoxifen. Initial gating is performed on the CD45^*−*^ CD31^*−*^ CD326^*−*^ Mcam^*−*^ Pdgfra^+^ population. Adventitial fibroblasts are Sca1^+^ and the Sca1^*−*^ population is made up of the alveolar fibroblast. After sorting cells were seeded for 48 hours on tissue culture plastic with either 10 ng/ml TGF*β*, 500 nM PGF2*α*, or media control. (B) Gene expression analysis via qPCR in untreated adventitial and alveolar fibroblasts quantifying the expression of population specific marker genes confirms the identity of target fibroblasts. (C) Quantification of transitional cluster maker genes after 48-hour challenge demonstrates the potential for FPr to induce the transitional state in adventitial fibroblasts. This is not observed in alveolar fibroblasts or in adventitial fibroblasts lacking the FPr, (D) Quantification of fibrotic cluster marker genes after 48-hour challenge confirms that TGF*β* promotes entry of both adventitial and alveolar fibroblasts into the fibrotic state independent of FPr status. Statistical significance between groups denoted by * : *p < 0.05 ** p<0.005 *** p<0.0005. Statistical significance between treatment and media control denoted by & : & p< 0.05

## DISCUSSION

Idiopathic pulmonary fibrosis remains a clinical challenge as an unmet need persists for well-tolerated and effective IPF therapeutics. While the fibroblast is recognized as a key lung effector cell population in IPF responsible for the synthesis and maintenance of extracellular matrix during aberrant injury repair, the emerging complexity of its biology including the functional importance of recently identified diverse fibroblast subsets has created new challenges for drug discovery. The bioactive eicosanoid, prostaglandin F2*α*, acting through its cognate receptor FPr has been implicated as a facilitator of fibrogenesis in IPF (32), however, a detailed understanding of the target mesenchymal population(s) influenced by PGF2*α* in IPF is lacking. Thus, to assess the role of PGF2*α*/ FPr signaling in IPF mechanistically, we utilized both genetic and pharmacologic approaches in two mouse models of IPF to affirm a contribution of FPr signaling to lung fibrogenesis. This also established an effect size for this pathway equivalent, but not additive, to that observed with the clinical antifibrotic Nintedanib. Then, using unbiased single cell RNA sequencing and *in vitro* validation, we localized *Ptgfr* expression predominantly within an adventitial fibroblast subpopulation which was capable of being selectively reprogrammed to a recently described “inflammatory/transitional” cell state in a PGF2*α* dependent manner. Collectively, our findings support a role for PGF2*α* signaling in IPF and mechanistically identify a target fibroblast subpopulation while also establishing a benchmark for disruption of this pathway in mitigating fibrotic lung disease.

For this study, the role of FPr signaling was first assessed in a clinically relevant model of spontaneous pulmonary fibrosis, the I^*ER*^-*Sftpc*^*I*73*T*^ mouse, which we have shown to recapitulate many characteristic features of IPF including histopathology, restrictive physiology, and biomarkers also found in human IPF. The translational relevance of this model was further supported by demonstration of the potential for nintedanib to rescue partially the fibrotic phenotype of the *Sftpc*^*I*73*T*^ mouse (**Figure S3**), in line with both other preclinical models and the observed clinical experience (8, 9, 30). Leveraging this model in combination with concomitant genetic ablation of FPr (**Figure S1**), we found that disruption of PGF2*α* signalling conferred a survival advantage and attenuated the fibrotic burden similar in magnitude to that of Nintedanib alone (**Figures 1**, **2, S3**) here or in the bleomycin-challenged FPr knockout mouse (32). We then extended these findings using 2 pharmacologic inhibitors of FPr signaling administered in an intervention protocol in the bleomycin mouse model which permitted the temporal segregation of early injury/inflammation from late fibrosis (**Figure S4**). Both OBE022, an FPr antagonist in clinical development as a tocolytic (35, 36), and BAY-6672, a quinolone based FPr antagonist shown to attenuate silica induced lung fibrosis in mice (37), each blocked fibrotic endpoints with effect sizes similar to Nintedanib alone. Importantly, a combination strategy of Nintedanib with either genetic or pharmacological targeting of FPr signaling was not additive or synergistic for any fibrotic endpoint (**Figure 3; Figure S4**). While this was surprising given that the known triple receptor kinase targets of Nintedanib are distinct from PGF2*α*/FPr signaling, our findings do not exclude that these two pathways can intersect at the same profibrotic cell population. Given current clinical trial design for potential IPF therapeutics, development of new first-in-class antifibrotics may not be as straightforward as simply targeting molecular pathways distinct from existing clinical therapies.

Despite the plethora of established and emerging targets for IPF, nearly all *in vitro* and *in vivo* approaches continue to measure efficacy based on the inhibition of the fibroproliferative state of the bulk lung mesenchyme (11, 55). With the rapid dissemination of single cell transcriptomic technology, a refined understanding of the spatial localization and pathological behavior of fibroblasts throughout the aberrant remodeling in both human IPF and murine fibrosis platforms is emerging (19-23, 53). Specifically, from computational inference analysis, lineage tracing, and *in vitro* modeling, anywhere from 6-9 mesenchymal cell subtypes have now been described with pathological collagen-producing fibroblasts (*Cthrc1*^+^ or myofibroblast). These may arise from several of these lineages within spatially distinct compartments of the lung including the *Col13a1*^+^,*/ Npnt*^+^ alveolar (lipo-) fibroblast of the proximal alveolar space as well as *Col14a1*^*H*^*i* / *Pi16*^+^ mesenchymal cells found in the lung adventitia although the predominant cell of origin remains a point of contention (20, 56). Embedded in several of these studies is the important observation that under a variety of exogenous stimuli (cytokines, bleomycin injury) the trajectory to a “fibrotic” MC population may involve prior entry and exit from a profibrotic intermediate state conventionally designated as “transitional” or “inflammatory” (22, 23).

Using our *Sftpc*^*I*73*T*^ fibrosis model, ScRNAseq analysis supports and extends these observations by both affirming the identity of all previously identified MC populations (**Figure 5**) and establishing a fibrotic trajectory for both alveolar and adventitial fibroblast populations to pathological fibroblasts through an inflammatory / transition state in the absence of an exogenous injury (e.g. bleomycin). RNA velocity also revealed that only the trajectory of the adventitial population was disrupted by deletion of FPr (**Figure 6A**). Importantly, FPr deletion did not alter the early inflammatory response (**Figure 3**) supporting a mechanism of action at the level of the fibroblast. The role for FPr signaling in fibroblast reprogramming events was further supported by the finding that in the *Sftpc*^*I*73*T*^ murine lung *Ptgfr* expression occurs predominantly within the mesenchyme (**Figure 5C**) with the highest levels spatially restricted mainly to *Col14a1*^*High*^/*Pi16*^+^ adventitial fibroblasts. A minor component is found in the alveolar population (**Figure 5C**). Interestingly, *Col14a1*^*High*^/*Pi16*^+^ adventitial MCs, which may also represent the murine homologue of the previously identified *Has1*^+^ mesenchymal populations described in human IPF (17), were the sole fibroblast population that maintained *Ptgfr* expression throughout the fibrotic remodeling following induction of mutant *Sftpc*^*I*73*T*^ (**Figure S9**) raising the potential that PGF2*α* signaling could modulate all or part of this trajectory in this population. This was confirmed using a reductionist approach isolating each of these populations and showing *in vitro* that it is the adventitial and not the alveolar population that demonstrates FPr dependence for entry into the transitional/inflammatory state (**Figure 7**).

The granular analysis of fibroblast subpopulations and their trajectories viewed via single cell analysis combined with our *in vitro* observation that FPr signaling selectively induces the transitional/inflammatory state only in the adventitial fibroblast, it appears likely that a profibrotic (*Cthrc1/Col1a1*^*high*^) population can be derived from at least two precursor mesenchymal cell populations via a common intermediate state with entry governed by a variety of cues. As shown in **Figure 8**, for the adventitial fibroblast, PGF2*α* represents at least one key driver of entry into a transitional/ inflammatory state. We speculate that one of more of the signaling kinases inhibited by Nintedanib would spatially overlap in the same adventitial population with resultant equivalency of effect size seen by their respective inhibition. This model does not preclude and, in fact, may support alternative fibroblasts, including the alveolar population, from generating profibrotic (*Cthrc1*) fibroblasts by arriving at the terminal profibrotic state through a separate signaling cascade (such as IL-1*β*) via the same or a parallel transitional/ inflammatory state. It also provides a plausible explanation for why fibrogenesis is only partially attenuated as there likely exists a large number of post-injury, pro-remodeling signals that each contribute to the development of the fibrotic phenotype through one or more of these precursor populations. Intervention in one or more of these signals may slow the development of a terminal fibrotic fibroblast population from a single compartment but does not comprehensively capture all pathways across multiple compartments that simultaneously influence human IPF.

**Figure 8.**
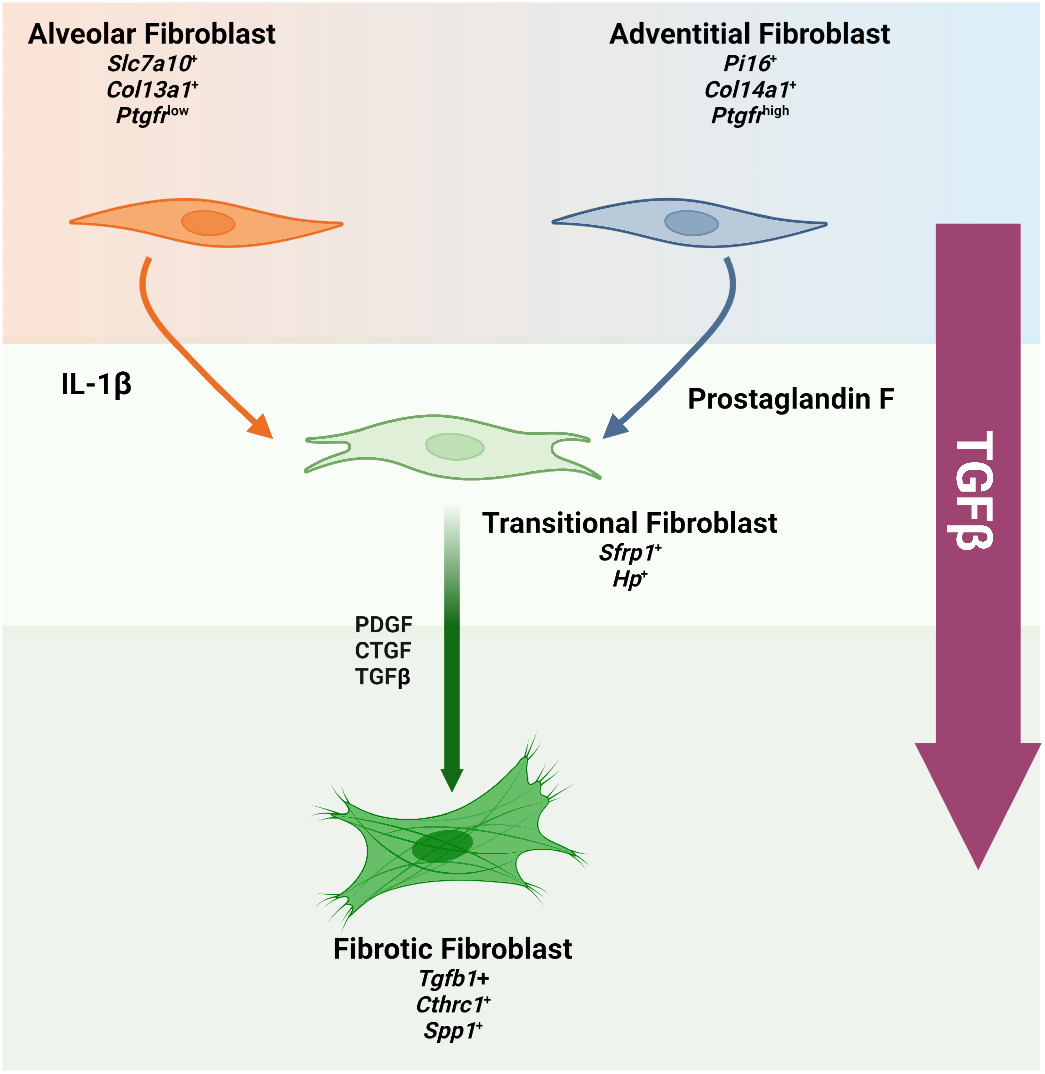
Summary of Potential Role for Prostaglandin (PG) F2α as a Driver of Fibroblast Heterogeneity in the Fibrotic Lung. Through a combination of single cell and *in vitro* validation we further define TGF*β* signaling as a driver of terminal fibroblast activation. Independent of adventitial or alveolar origin, TGF*β* is a classical, potent profibrotic ligand. Alternatively, within the fibrotic lung milieu exists several previously described alternative drivers of fibroblast heterogeneity including Il-1*β*, PDGF, and CTGF. Adding to this list of drivers we identify PGF2*α* and a novel, adventitial specific diver of the transitional state. Once in this state, the fibroblast may be more sensitive to TGF*β* signaling and accelerate the expansion of the terminal fibrotic fibroblast population.

We also note that while PGF2*α* promoted development of the transition state it did not, in isolation, increase markers of the terminal profibrotic state. We also found that addition of TGF*β* to either alveolar or adventitial fibroblasts in culture successfully drove each of these cells to a profibrotic state while decreasing expression of markers of the transitional / inflammatory state (**Figure 7C**). The role of TGF*β* as a potent profibrotic signal has been well documented and is a significant area of interest for human clinical trials. In this study we observed that TGF*β* expressing fibroblasts clustered into the terminal fibrotic population and we have also shown previously that there is also a significant amount of TGF*β* in the fibrotic milieu arising from the immune compartment entering the profibrotic remodeling phase of injury resolution (34). This potent signal, arising from multiple sources, may represent a “system override” that increases the size of the fibrotic populations observed in our model by pushing cells rapidly through the transitional/ intermediate state or present a second “hit” profibrotic signal to cells in the transitional state. Our *in vitro* data demonstrating the efficacy of TGF*β* to induce the fibrotic state independent of an initial FPr signal in both adventitial and alveolar derived fibroblasts further reinforce the potential of this molecule to act independently of an upstream pathway driven by PGF2*α*. This finding contrasts with the study by Oga *et at*. (32) where both PGF2*α* and TGF*β* were each shown to increase collagen production and enhance proliferation in cultures of bulk isolated fibroblasts. We speculate that, in that model system, PGF2*α* mediated entry of the adventitial subpopulation into the transitional state results in a transitional state-dependent stimulation of a second fibroblast population that was not present in our purified populations. Confirmational studies of the trajectory and transcriptomic dynamics of the fibroblast populations will require the generation of cell population specific reagents for lineage tracing as well as further genetic and/or pharmacologic interrogation of other pathways. In conclusion, using the combined application of IPF genetic models and single cell technology to address the fibroblast heterogeneity arising throughout the aberrant injury/repair process observed with pulmonary fibrosis, we have established a role for PGF2*α* signaling in PF acting through a key mesenchymal population, the adventitial fibroblast. Our work here begins to elucidate the importance of the transitional profibrotic state that multiple fibroblast subpopulations enter and highlights the relevance to fibrotic remodeling when targeted through intervention strategies.

## AUTHOR CONTRIBUTIONS

MFB and GAF developed the concept. MFB, LR, ST, TC, and GAF designed the experiments. ST and YT performed *in vivo* animal experiments; LR, ST, SI, KC, TD, and AM performed in vitro experiments and endpoint analyses for *in vivo* studies; UT and SG performed mass spectrometric analyses of eicosanoids; WRB and CC performed bioinformatic analysis. AM performed flow cytometry; TC, ST, AM, LR, and MFB analyzed data, generated figures, and interpreted results; LR and MFB drafted manuscript; GAF, MFB, and LR edited the manuscript. All authors reviewed and approved the final version prior to submission.

## ACKNOWLEDGMENTS

Michael F. Beers is an Albert M. Rose Established Investigator of the Pulmonary Fibrosis Foundation and is the Robert L. Mayock and David A. Cooper Professor of Medicine. Garret A. FitzGerald holds a Merit Award from the American Heart Association and is the McNeil Professor of Translational Medicine and Therapeutics. We thank Jeremy Katzen for helpful discussions and the PENN-CHOP Lung Biology Institute Informatics team for guidance.

