## Supplemental Data for "Disruption of Prostaglandin F_2α_ Receptor Signaling Attenuates Fibrotic Remodeling and Alters Fibroblast Population Dynamics in A Preclinical Murine Model of Idiopathic Pulmonary Fibrosis"

### SUPPLEMENTAL METHODS

#### *$Sftpc^{I73T}$ Targeting Vector and Recombineering strategy for generation of SP-$C^{I73T}$ founder line*

Homozygous FPr knockout mice have been previously described (32) and were kindly provided by Shuh Narumiya (Kyoto University Faculty of Medicine, Japan). The breeding scheme to generate triple homozygous is detailed in *Supplemental Methods*. All mouse strains and genotypes generated for these studies were congenic with C57BL/6. Both male and female animals (aged 8-14 weeks) were utilized in tamoxifen induction protocols. All mice were housed under pathogen free conditions in an AALAC approved barrier facility at the Perelman School of Medicine, University of Pennsylvania. All experiments were approved by the Institutional Animal Care and Use Committee at the University of Pennsylvania.

#### *Generation of F Prostanoid receptor (FPr)- deficient $I^{ER}$-$Sftpc^{I73T}$ mouse model s*

Triple homozygous ($I^{ER}$-SP-$C^{I73T/I73T}$ $FlpO^{+/+}$/ $Fpr^{-/-}$) mice lacking *Ptgfr* were generated by initially crossing male $Sftpc^{I73T/I73T}$ / $Flp^{+/+}$ mice with female $FPr^{+/-}$ mice. The presence of SP-$C^{I73T}$, FlpO, and FPr alleles were assessed by PCR analyses. Multiple intercrosses resulted in generation of $Sftpc^{I73T/I73T}$/$Flp^{+/+}$/$Ptgfr^{-/-}$ ("FPr knockouts"), $Sftpc^{I73T/I73T}$/$Flp^{+/+}$/$Ptgfr^{+/-}$ ("FPr heterozygotes") and $Sftpc^{I73T/I73T}$/$Flp^{+/+}$/$Ptgfr^{+/+}$ ("FPr controls"). Additional controls included a FPr deficient, Sftpc wild-type strain ($I^{ER}$-$Sftpc^{WT/WT}$/$Flp^{+/+}$/$Ptgfr^{-/-}$).

#### *PCR based genotyping of mice*

DNA was extracted from tail snips obtained from pre-weaned mouse pups and then used as template for detection of the following alleles by polymerase chain reaction (PCR): $Sftpc^{WT}$, $Sftpc^{I73T}$, and $Sftpc^{I73T-Neo}$ - Multiplex PCR was done using a common reverse primer 3' downstream of the PGK-Neo insertion site, with alternate forward priming sites located upstream of or within the PGK-Neo cassette. Amplification was performed using Platinum Taq (Invitrogen), with the following primer sequences:
SP-C Fwd: TCACCCCTGTCCTCTCTGTC
PGK Fwd: TGGATGTGGAATGTGTGCGA
SP-C Rev: CCCAACTACATGGTGGTGCTA
The thermal cycling conditions were: 95°C for 3 minutes; 37 cycles of 95°C (30sec), 64°C (30 sec), 72°C (60 sec). For $Sftpc^{WT}$ this results in amplification of a 433 BP product. For $Sftpc^{I73T}$ alleles post-excision, 113 base pairs (including the FRT and HA sites) are added, resulting in a 446 bp band. For alleles pre-excision ($Sftpc^{I73T-Neo}$), an alternate forward priming site within the PGK promoter permits preferential amplification of a 271 bp band. R26Flp-O ER - Multiplex PCR was done using a common forward primer and two reverse primers (as described by Jackson Laboratories, Inc.). Amplification was performed using Platinum Taq (Invitrogen), with the following primer sequences:
Common Forward (oIMR8545): AAAGTCGCTCTGAGTTGTTAT
I73T Mutant Reverse (10507): TTATGTAACGCGGAACTCCA
Wild type Reverse (oIMR8546): GGAGCGGGAGAAATGGATATG
The thermal cycling conditions used were: 95°C for 3 minutes; 10 cycles of 95°C (30sec) 65 - 0.5 °C/cycle (30sec), 68°C (60sec); 28 cycles of 95°C(30 sec), 60°C (30sec) 72°C (60 sec). These conditions result in amplification of a 603 bp band (Flp-O negative) and a 309 bp band (Flp-O positive). *Ptgfr* Status - The thermal cycling conditions were: 95°C for 5 minutes; 35 cycles of 94°C (30sec), 65°C (30 sec), 75°C (60 sec). For $Ptgfr^{WT}$ this results in amplification of a 700 BP product. For $Ptgfr^{KO}$ alleles this results in a 450 bp band. Amplification was performed using the following primers:
FP1: GCC CAT CCT TGG ACA CCG AGA TTA TCA A-3'
FP2: AGA GTC GGC AAG CTG TGA CTT CGT CTT-3'
FP3: TGA TAT TGC TGA AGA GCT TGG CGG CGA A-3'

#### *Determination of FPr inhibitor Efficacy in A Bleomycin Induced Mouse Lung Fibrosis Model*

OBE-022 and BAY6672 were evaluated in a bleomycin (BLM) induced lung fibrosis model performed by HD Biosciences, Ltd. Shanghai, PR China. For each compound, 85 male C57/Bl6 male mice were randomly divided into 5 groups: 1) Group 1 (17 mice): Sham group, mice were administered with PBS (i.t) and received vehicle (0.5% [w/v] HPMC and 0.02% [v/v] Polysorbate 80) in Water (p.o., QD); 2) Group 2 (17 mice): Model group, mice were administered with BLM (0.66mg/kg, i.t) and received vehicle (0.5% [w/v] HPMC and 0.02% [v/v] Polysorbate 80) in Water (p.o., QD); 3) Group 3 (17 mice): OBE022 (100mpk) or BAY6672 (30 or 100 mpk). Mice were administered with BLM (0.66mg/kg, i.t) and received OBE022 or (100mpk, p.o., BID or BAY6872 (30 or 100 mpk)); 4) Group 4 (17 mice): Calico-006, 300mpk, mice were administered with BLM (0.66mg/kg, i.t) and received Calico-006 (300mpk, p.o., BID); 5) Group 5(17 mice): Nintedanib 60mpk, mice were administered with BLM (0.66mg/kg, i.t) and received Nintedanib (60mpk, p.o., QD). The mice were administered with BLM (0.66mg/kg, i.t.) on day 1 and randomized the mice and dosed compounds from day 5 after BLM injection. Compound treatment occurred for 17 days of an entire study period lasting 21 days. End point assays included (i) Total body weight; (ii) BALF cell counts and cytological differentials; BALF soluble collagen content using Sircol assay (ii) Histopathological examination for collagen deposition and inflammation quantification (iv) IHC analysis (staining and quantification) of $\alpha$-SMA. Histopathological evaluation of lung using Masson Trichrome Staining and fibrosis was evaluated by Modified Ashcroft scoring. Whole slide scanning and image analysis was performed to quantify $\alpha$-SMA in the fibrotic lung tissues by positive area.

### SUPPLEMENTAL FIGURES

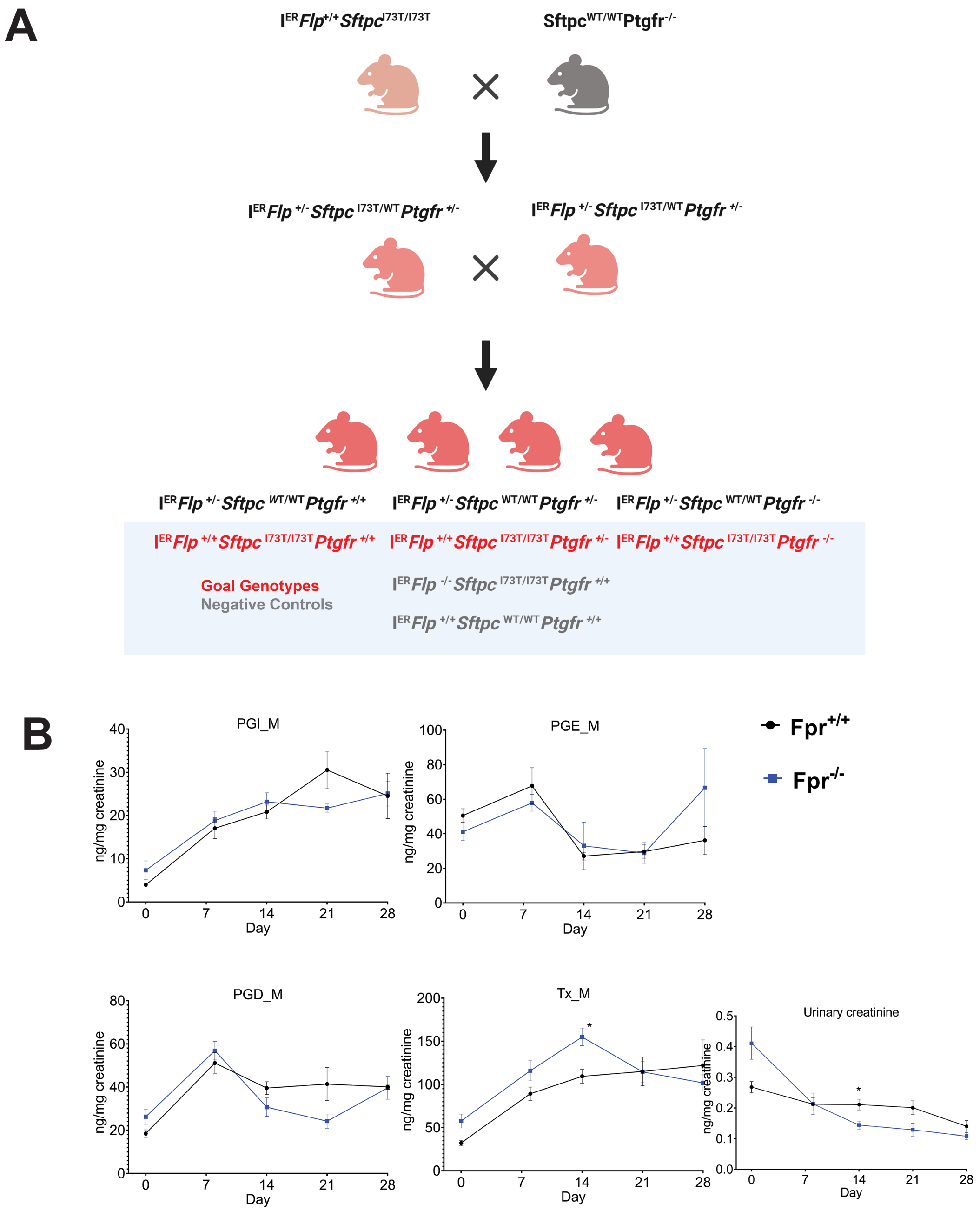


**Figure S1** *Deletion of Ptgfr reduces morbidity and mortality in IER-SftpcI73T mice.* (A)*Ptgfr* mRNA content of whole lung mRNA isolated from generated lines of I$^{ER}$-*Sftpc*$^{I73T}$/*Ptgfr* mice deficient in 0, 1, 2 *Ptgfr* alleles assayed by qRT-PCR; (B) Schematic of single, split ip or split OG dosing strategy employed for Tamoxifen induction of I$^{ER}$-*Sftpc*$^{I73T}$/*Ptgfr*$^{-/-}$ and I$^{ER}$-*Sftpc*$^{I73T}$/*Ptgfr*$^{+/+}$ cohorts; (C) Representative weight loss curve from a single cohort containing I$^{ER}$-*Sftpc*$^{I73T}$/*Ptgfr*$^{-/-}$ (n=20) and I$^{ER}$-*Sftpc*$^{I73T}$/*Ptgfr*$^{+/+}$(n=13) controls * P < 0.05 vs controls(D) Aggregate Kaplan Meier curve for I$^{ER}$-*Sftpc*$^{I73T}$/*Ptgfr*$^{-/-}$ mice from 3 cohorts separately induced with either single ip or split ip doses of Tamoxifen in Corn Oil with total numbers of each Ptgfr genotype shown. Negative Controls consisted of *Sftpc*$^{W}$*T* C57BL/6 mice given Tamoxifen or uninduced I$^{ER}$-*Sftpc*$^{I73T}$/*Ptgfr*$^{-/-}$ animals. p values versus I$^{ER}$-*Sftpc*$^{I73T}$/*Ptgfr*$^{-/-}$ obtained by log-rank testing are shown.

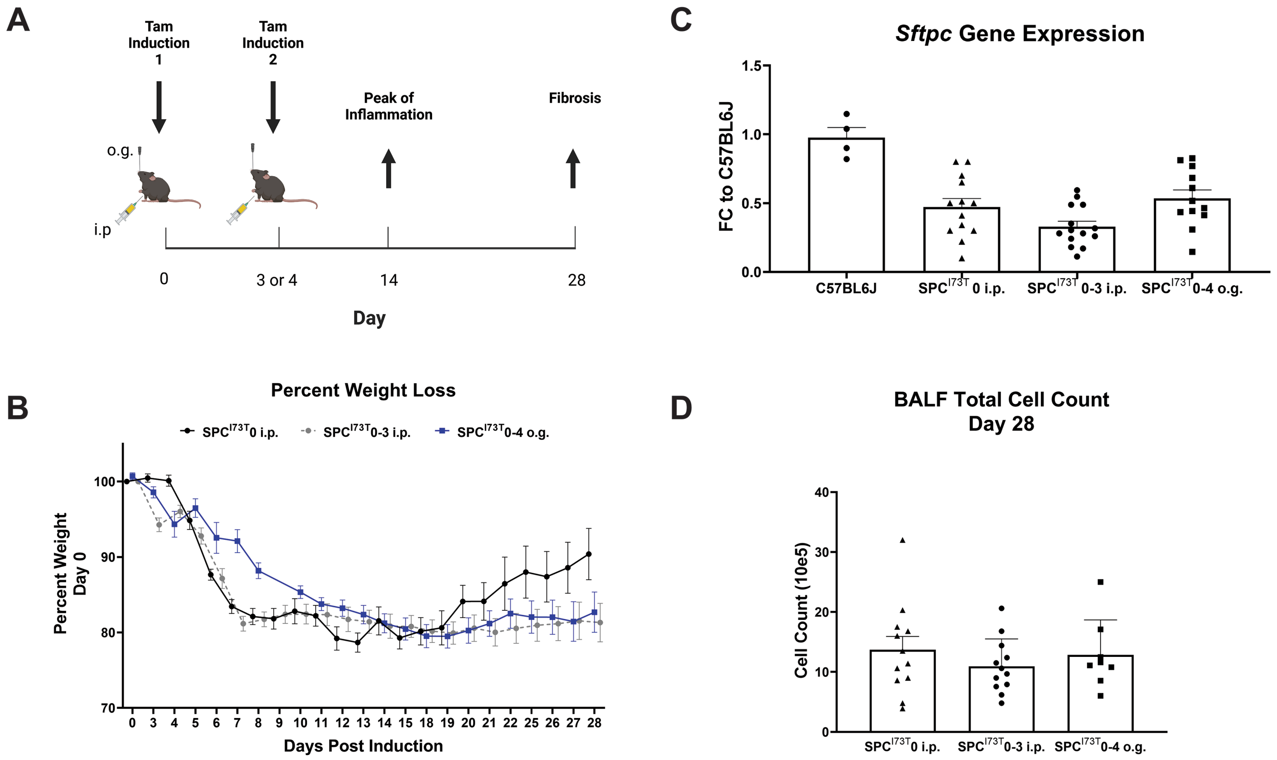


**Figure S2** *Alternative Protocols for tamoxifen in Sftpc$^{I73T}$ do not alter model outcomes.* (A) Schematic describing three approaches to administer tamoxifen in I$^{ER}$-*Sftpc*$^{I73T}$ mice: Single intraperitoneal delivery (n=13) , split intraperitoneal delivery on day 0 and 3 (n=14), or split gavage on day 0 and 4 (n=12). (B) Weight loss nadir in I$^{ER}$-*Sftpc*$^{I73T}$mice is similar across all administration protocols with a more gradual progression observed using the split dose oral gavage protocol. (B) Expression of Sftpc as measured by qPCR is comparable in all three protocols. (D) Total cell count in BALF 28 days after first tamoxifen induction is comparable across all three protocols.

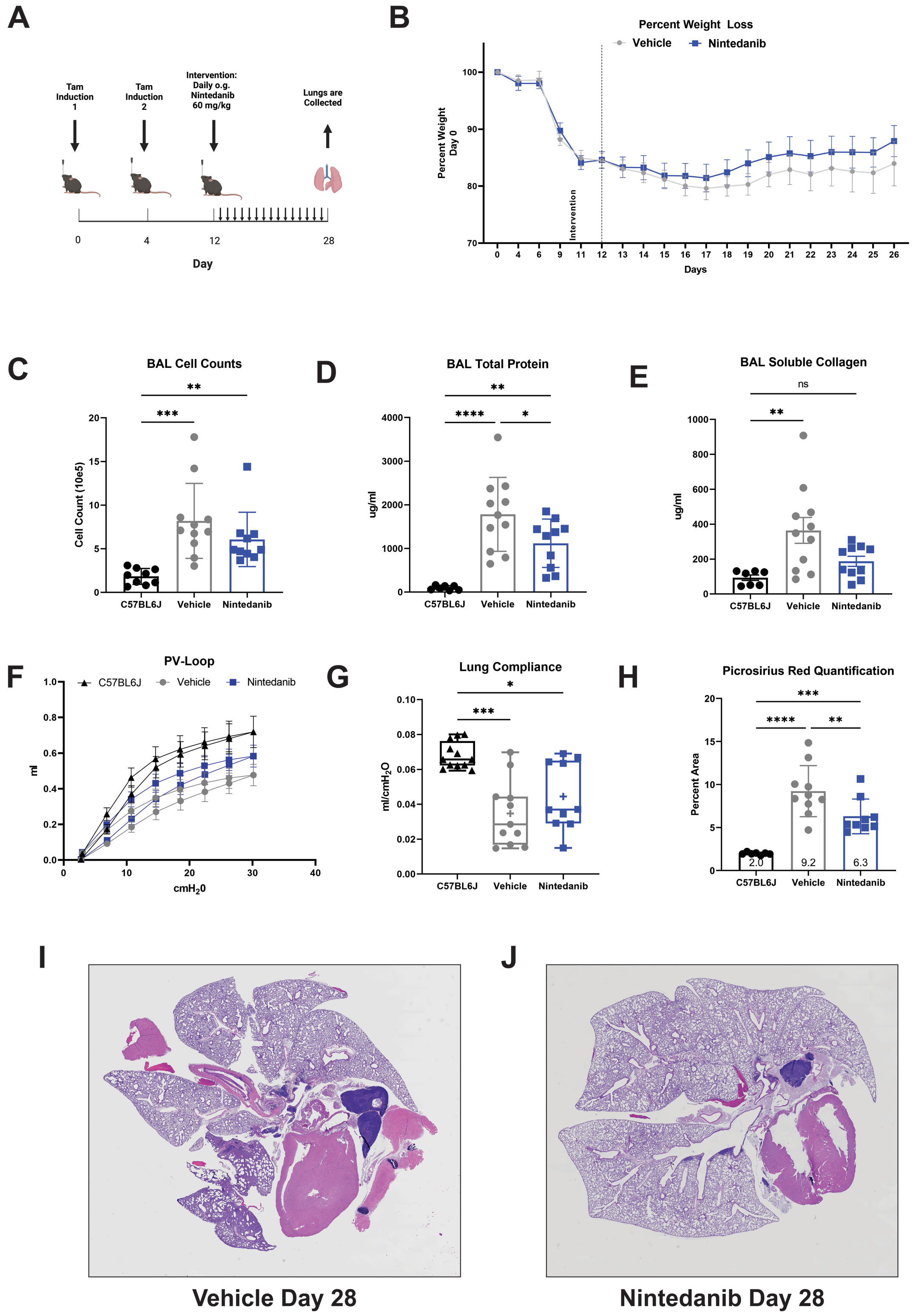


**Figure S3** *Effect of nintedanib treatment on spontaneous fibrotic lung remodeling following oral tamoxifen induction of Sftpc$^{I73T}$ expression.* (A) Schematic of protocol used for tamoxifen induction of *Sftpc*$^{I73T}$ mice and nintedanib treatment. (B) Weight loss in induced *Sftpc*$^{I73T}$ mice receiving nintedanib (n=13) or vehicle (n=13) at Day 12. (C-E) BALF collected from *Sftpc*$^{I73T}$ mice 28 days post tamoxifen induction demonstrates increased cell count, total protein, and soluble collagen (Sircol™ Fibrillar collagen assay) as compared to C57BL/6 (n=7) controls. Nintedanib partially mitigates these metrics with a significant decrease in BALF total protein after intervention. (F-G) Lung compliance is significantly reduced in *Sftpc*$^{I73T}$ mice with a nonsignificant partial mitigation observed after nintedanib treatment. (H) Quantification of fibular collagen in histological sections (reported as % PSR Stained Area) from *Sftpc*$^{I73T}$ lungs 28 days post-tamoxifen demonstrates a significant reduction in collagen deposition after nintedanib intervention. (I-J) Representative histology from I$^{ER}$-*Sftpc*$^{I73T}$ mice 28 days post tamoxifen induction.

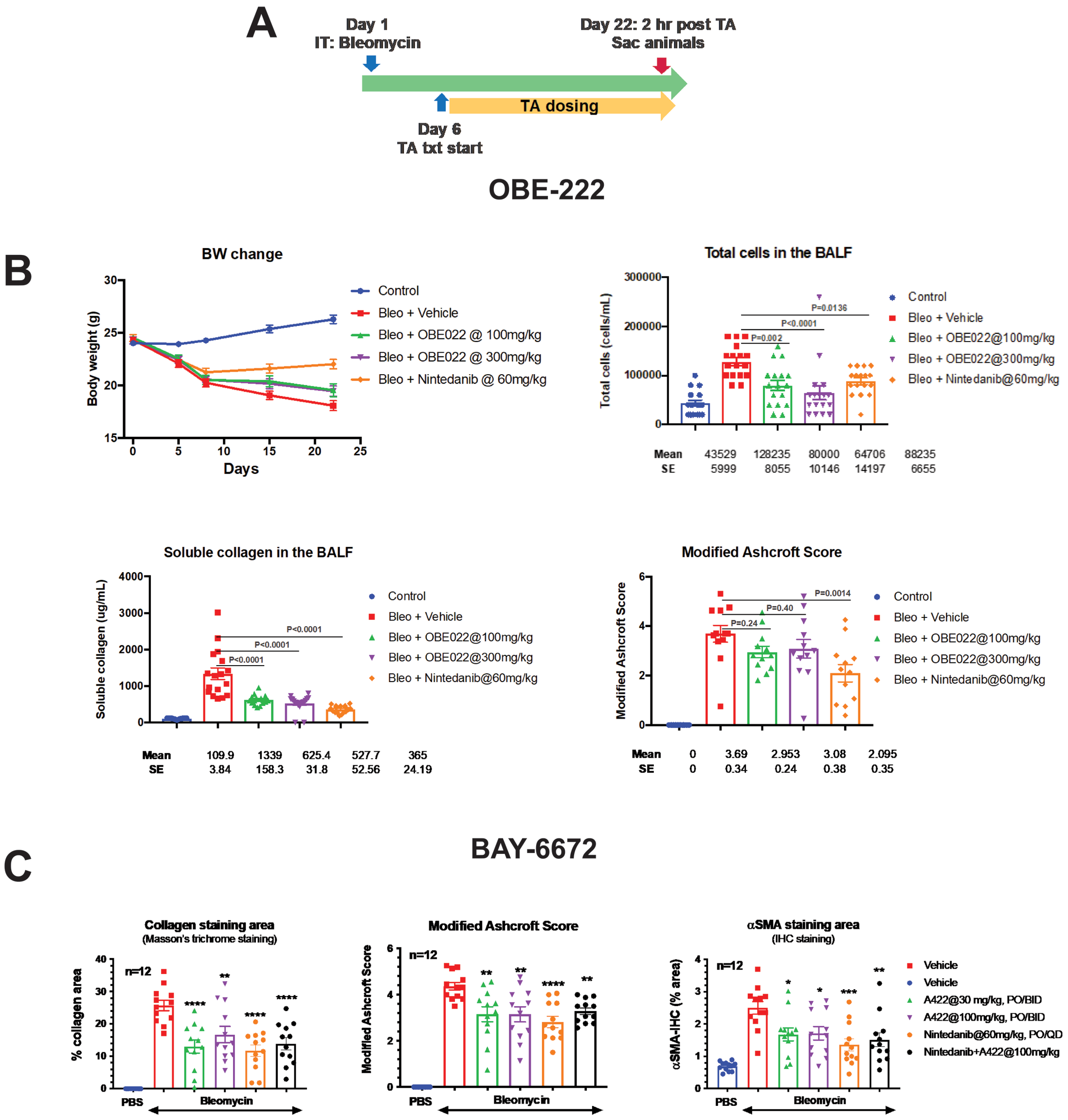


**Figure S4** *Pharmacological Inhibition of FPr Signaling Decreases Fibrotic Endpoints After Bleomycin Injury.* (A) Schematic of pharmacological intervention strategy used for OBE-222 and BAY-6672. Treatment was started on Day 6 and maintained through day 22 with final collection occurring 2 hours post final dose administration. (B) Endpoint analysis of OBE-022 treatment compared to nintedanib intervention. All groups were made up of n=17 animals. No significant differences were observed in body weight across all interventions. Cell count and soluble collage in BALF after bleomycin challenge was significantly reduced in all intervention protocols as compared to vehicle controls. Histological scoring applying modified Ashcroft Score resulted in a significantly lower score after nintedanib intervention. (C) Histological scoring of BAY-6672 (alternative identifier A422), nintedanib, and combined interventions. All groups were made up of n=12 animals. Modified Ashcroft scoring demonstrates mitigation of fibrotic severity post bleomycin injury in all interventions with no significant difference when nintedanib is combined with BAY-6672. This trend is also observed in collagen and smooth muscle actin quantification.

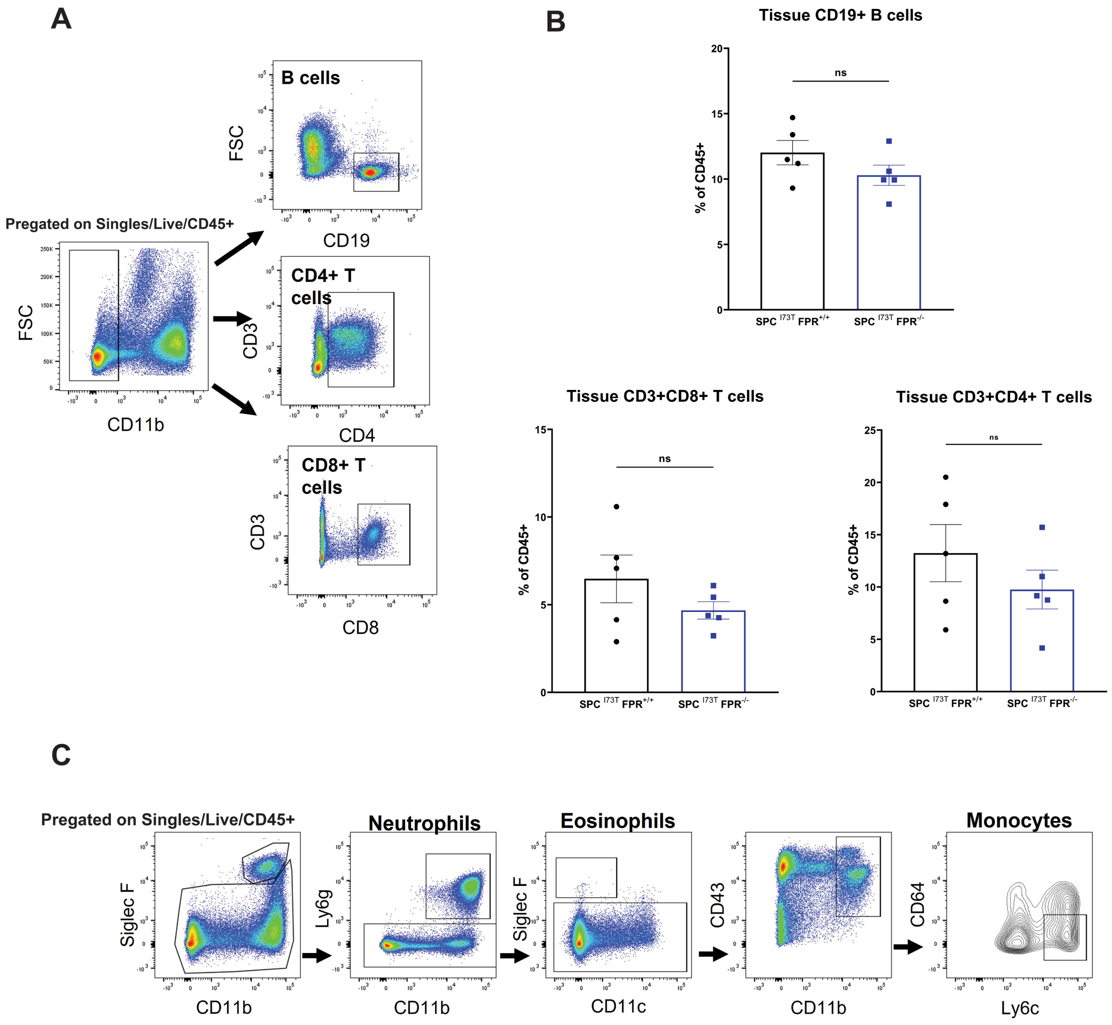


**Figure S5** *Ptgfr Deficiency Does not Alter Tissue Lymphocyte Populations in $I^{ER}$-$Sftpc^{I73T}$*.(A) Gating strategy used to quantify B-cells, CD4 T cells, and CD8 T cells in single cell suspensions derived from murine lungs. (B) No significant difference in tissue lymphocyte percentages was observed between I$^{ER}$-*Sftpc*$^{I73T}$/*Ptgfr*$^{+/+}$ (n=5) and I$^{ER}$-*Sftpc*$^{I73T}$/*Ptgfr*$^{-/-}$(n=5) mice (C) Gating strategy used to quantify neutrophils, eosinophils, and Ly6c hi monocytes in single cell suspensions derived from murine lungs.

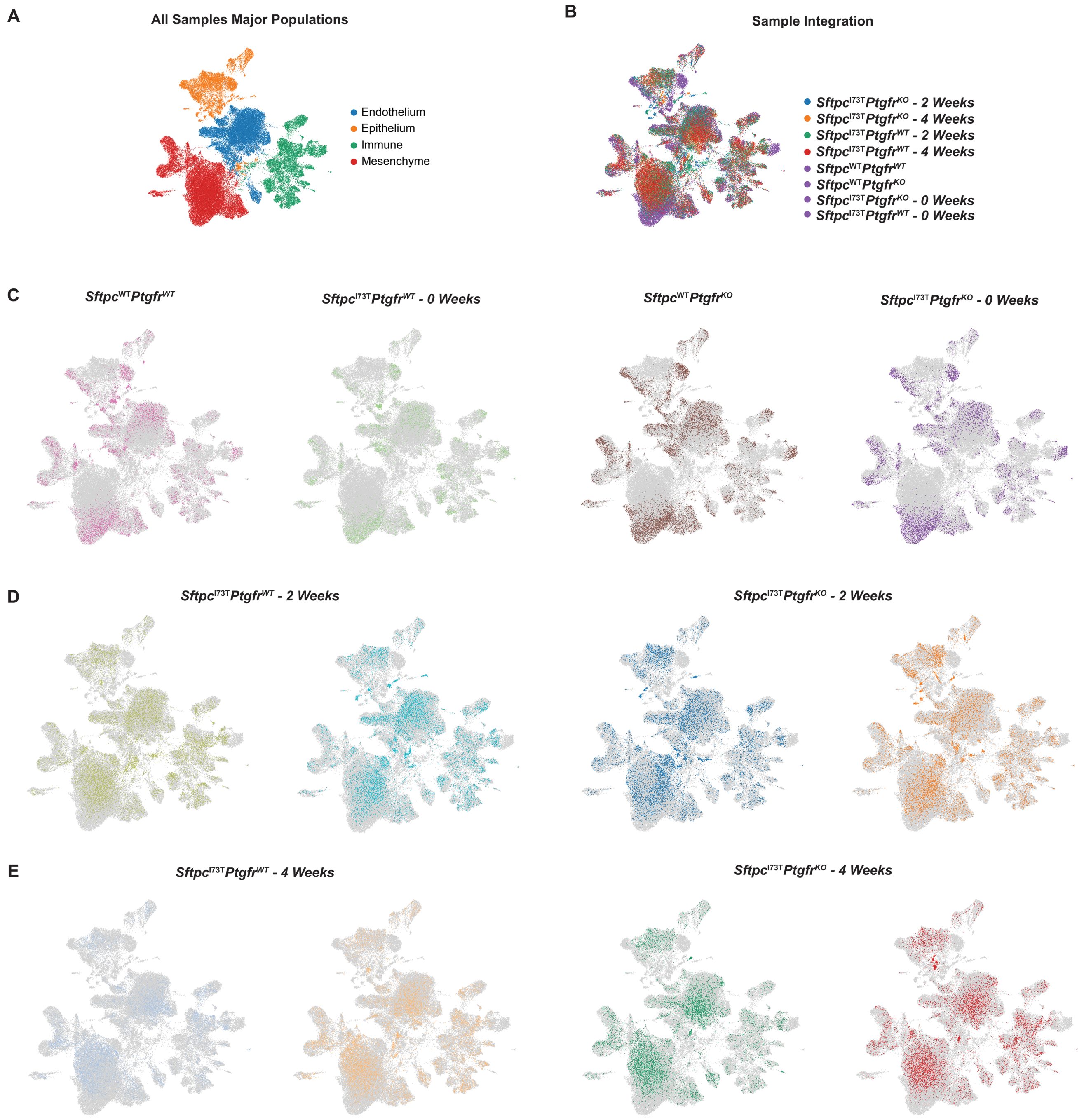


**Figure S6** *Single Cell Sample Integration in $I^{ER}$-$Sftpc^{I73T}$* (A) UMAP projection of integrated data set identifies the four major cell compartments in murine lungs (B) UMAP projection of integrated data set with color designations for each timepoint and genotype. (C-E) UMAP projections of integrated divided by individual samples with single color designations.

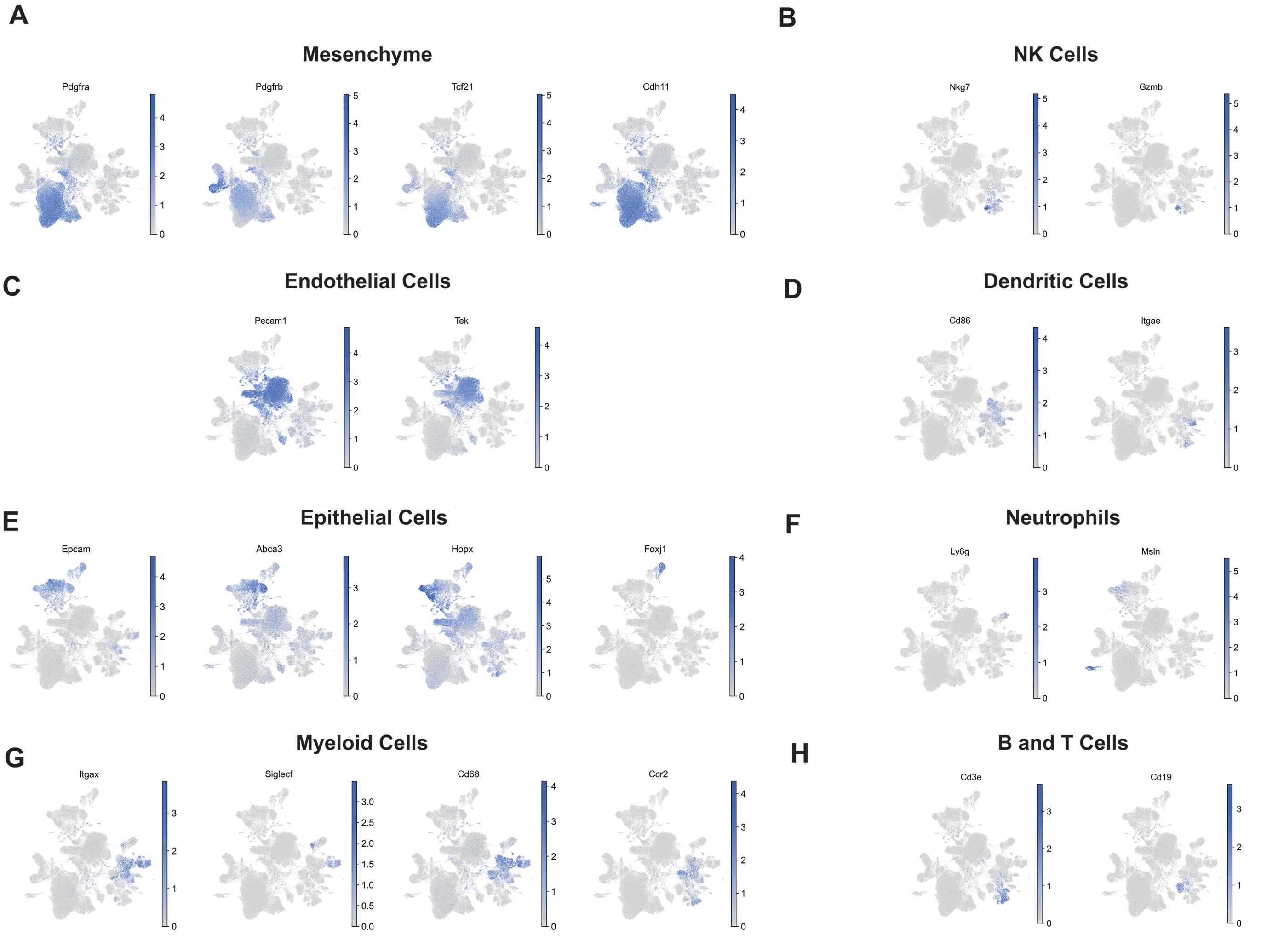


**Figure S7** *Cell Type Identification in Integrated $I^{ER}$-$Sftpc^{I73T}$ Single Cell Data* . (A)UMAP projections of integrated data sets highlighting cluster genes used to identify mesenchymal subsets. (B) UMAP projections of integrated data sets highlighting cluster genes used to identify natural killer cells. (C) UMAP projections of integrated data sets highlighting cluster genes used to identify endothelial cells. (D) UMAP projections of integrated data sets highlighting cluster genes used to identify dendritic cells. (E) UMAP projections of integrated data sets highlighting cluster genes used to identify epithelial cells. (F) UMAP projections of integrated data sets highlighting cluster genes used to identify neutrophils. (G) UMAP projections of integrated data sets highlighting cluster genes used to identify myeloid cells. (H) UMAP projections of integrated data sets highlighting cluster genes used to identify lymphocytes.

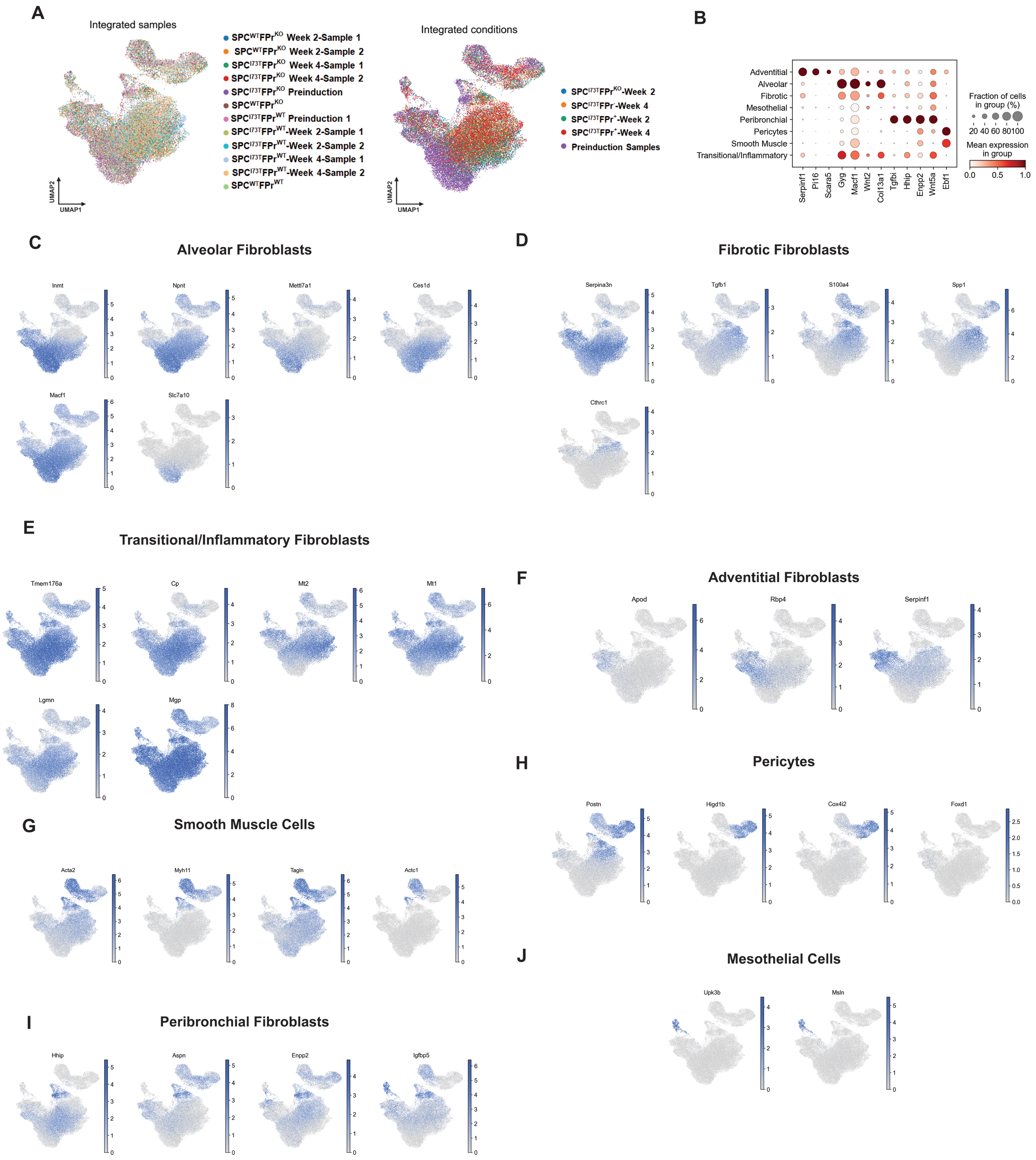


**Figure S8** *Mesenchymal Cell Cluster Identification in Sftpc$^{I73T}$ Integrated Single Cell Data Set.* (A) UMAP projection of integrated mesenchymal compartment with color designations for each timepoint and genotype. (B) Alternative Marker genes for mesenchymal subsets graphed as a gradient dot plot. (C) UMAP projections of integrated mesenchyme highlighting cluster genes used to identify alveolar fibroblasts. (D) UMAP projections of integrated mesenchyme highlighting cluster genes used to identify fibrotic fibroblasts. (E) UMAP projections of integrated mesenchyme highlighting cluster genes used to identify transitional/inflammatory fibroblasts. (F) UMAP projections of integrated mesenchyme highlighting cluster genes used to identify adventitial fibroblasts. (G) UMAP projections of integrated mesenchyme highlighting cluster genes used to identify smooth muscle cells. (H) UMAP projections of integrated mesenchyme highlighting cluster genes used to identify pericytes. (I) UMAP projections of integrated mesenchyme highlighting cluster genes used to identify peribronchial fibroblasts. (J) UMAP projections of integrated mesenchyme highlighting cluster genes used to identify mesothelial cells..

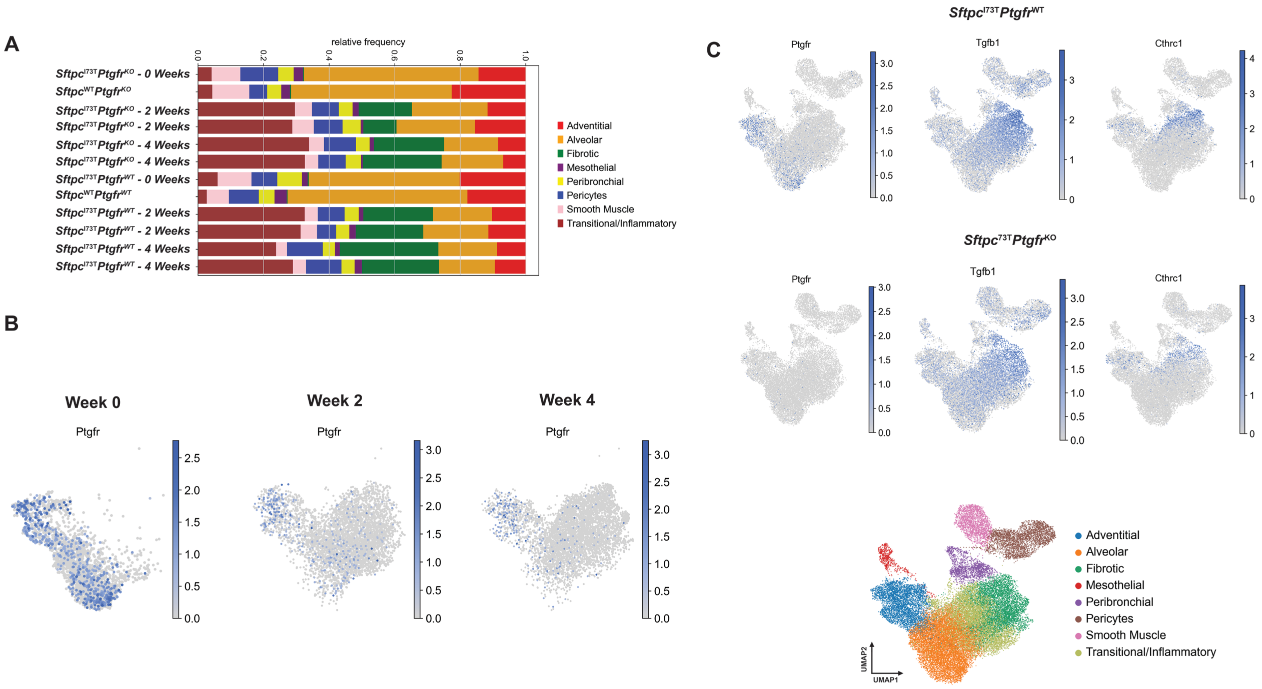


**Figure S9** (A) Frequency plot of mesenchymal subsets within individual samples. (B) UMAP projections of *Pdgfra* positive mesenchymal clusters divided by timepoint and highlighting the expression of *Ptgfr*. (C) UMAP projection of integrated mesenchymal clusters divided by genotype and highlighting profibrotic markers.
